# The importance of environmental parameters and mixing zone in shaping estuarine microbial communities along a freshwater-marine gradient

**DOI:** 10.1101/2022.03.07.483378

**Authors:** R.R.P. Da Silva, C.A. White, J.P. Bowman, L. Bodrossy, A. Bissett, A. Revill, R. Eriksen, D.J. Ross

## Abstract

Microbial communities are important elements in the marine environment, contributing to nutrient cycling and biogeochemical processes. Estuaries comprise environments exhibiting characteristics from freshwater to marine, leading to distinct microbial communities across this environmental gradient. Here, we examine the spatial dynamics of microbial communities in Macquarie Harbour, an estuarine system on the West coast of Tasmania, Australia. Water was sampled along the estuary to explore the structure and composition of the microbial communities using 16S/18S rRNA gene amplicon sequencing. Multivariate analyses showed environmental variables and community compositions varying along a longitudinal (river to adjacent ocean) gradient at the surface. In the harbour, differences in the microbial community were observed between surface (0-1 m) and intermediate depths (4.5-11 m depth). The results of differential abundance, network and Partial Least Square analyses suggest that Macquarie Harbour is a mixing zone, where the distributions of archaeal, bacterial and eukaryotic communities are influenced by oceanic and riverine inputs. Coupled with the natural characteristics of the Harbour, the heterotrophic component of this microbial communities inhabiting the surface and intermediate waters may play important roles in the nutrient cycle in the studied area. These results provide critical insights into the Macquarie Harbour environment and the importance of understanding the role of microbial communities for similar systems elsewhere.

## INTRODUCTION

Microbial communities are a key component of aquatic systems, playing fundamental roles in biological and geochemical processes (Azam and Malfatti, 2007; Jiao et al., 2014; Guidi et al., 2016). As coastal systems are the confluence of oceanic, freshwater and terrestrial systems, microbial community function and structure is largely dependent on hydrological processes that influence basic environmental parameters such as oxygen, temperature, salinity, and nutrient gradients, all of which control microbial distribution and composition (Parsons et al., 2015; Bartl et al., 2018; Varkey et al., 2018; Chenard et al., 2019). Furthermore, environmental perturbations caused by climate change and anthropogenic activities have been reported as drivers of microbial dynamics (Jeffries et al., 2015; Cavicchioli et al., 2019). Therefore, disturbances in coastal systems influenced by both natural phenomena, as well as anthropogenic activities, can produce complex effects on microbial communities.

Continual high levels of nutrient inputs to coastal areas, particularly nitrogen and phosphorous, often have deleterious environmental consequences, such as persistent hypoxia (Howarth et al., 2011; Reed and Harrison, 2016). Hypoxia is considered to occur when dissolved oxygen (DO) levels fall below 2 mg L^-1^ (~62.5 μM) (Gray et al., 2002; Rabalais, 2002). Factors such as water column stratification and high rates of organic matter decomposition, which often occur in sheltered coastal areas, can facilitate persistent hypoxia (Rabalais, 2002; Diaz and Rosenberg, 2008; Thrash et al., 2017). Although hypoxia occurs naturally, an increase in its frequency and extent, coupled to anthropogenic activities, has been observed in coastal environments around the globe (Howarth et al., 2011; Wright et al., 2012; Liu et al., 2015; Jessen et al., 2017).

Macquarie Harbour is a semi-enclosed estuarine system located on the west coast of Tasmania (42°18’13.8”S; 145°22’22.7”E), Australia. It has a stratified water column and restricted oceanic water exchange. The harbour has fjord-like hydrodynamics, characterized by a brackish layer on top of denser, salty waters, with distinct bacterial communities inhabiting each water layer (Da Silva et al., 2021). Tannin rich freshwater, discharged by two major rivers, limits light penetration (< 2 m-3m), possibly limiting photosynthesis (Hartstein et al., 2016; Ross et al., 2016). Colder and saltier marine waters enter through a narrow and shallow entrance channel and sink, leading to the recharge and replenishment of bottom water dissolved oxygen (DO) (Cresswell et al., 1989; Hartstein et al., 2019). Below the halocline (> 10 m depth) DO concentrations are historically, naturally, low (4 - 6 mg O_2_ L^-1^), but in recent years, there has been a significant DO decline observed, with levels now 2 - 4 mg O_2_ L^-1^, and often lower at 25 m depth (Ross et al., 2021). This DO decline is thought to be driven by a combination of increasing sediment biological oxygen demand, possibly due to fish farm inputs, and a reduction in the scale and frequency of DO recharge events (Ross and Macleod, 2017). Biological oxygen demand in intermediate waters (i.e., oxic-anoxic transition zone) is also considered to be a significant driver of oxygen drawdown (Revill et al., 2016). This general trend toward lower DO has highlighted the need to better comprehend the drivers of water column oxygen dynamics.

Estuaries are areas where freshwater meets the marine environment, leading to distinct communities across environmental gradients, horizontally and/or vertically (Crump et al., 2004; Liu et al., 2014). Where stratification occurs each water layer can act as a barrier, controlling microbial distributions and associated biological processes (Agogue et al., 2011; Jing et al., 2015; Sunagawa et al., 2015; Damashek et al., 2016). In stratified environments, the availability of nutrients and energy substrates is key for determining and understanding microbial dynamics and their role in ecosystem function. While phytoplankton influence surface nitrogen concentrations, below the oxycline and in the absence of light, non-photosynthesizing bacteria will dominate biochemical processes, affecting nutrient and water column oxygen availability (Wakeham et al., 2007; Damashek and Francis, 2018; Wu et al., 2019). For instance, in tannin-rich environments, where light is limited below the surface, chemoautotrophic production (e.g., nitrogen fixation, nitrification, sulfur oxidation, methanogenesis) contributes to nutrient cycling within the oxic-anoxic transition layers (Taylor et al., 2001; Pimenov and Neretin, 2006; Arístegui et al., 2009; Santoro, 2009; Wu et al., 2019). Furthermore, riverine and other inputs, that lead to an increase in nutrient availability (e.g., ammonia: NH_4_^+^), suspended particulate matter (SPM), and turbidity, tend to favor heterotrophic bacteria over autotrophic phytoplankton (Ward, 2008; Smith et al., 2014; Damashek et al., 2016; Bartl et al., 2018). Therefore, enhanced heterotrophic bacterial activity is an important process that can increase oxygen consumption in vertically stratified estuaries, particularly those with limited light availability. As oxygen is the major source of energy for organic matter mineralization, declines in oxygen concentration can lead to shifts in organic matter metabolism with nitrogen, sulfur, and manganese cycling also common in oxygen-deplete systems (Berg et al, 2013; Colatriano et al, 2015; Bush et al, 2017; Liu et al, 2020). A better understanding of the relationship between environmental parameters and prevailing microbial communities will provide insight into population dynamics, as well as allowing investigation into potential drivers of oxygen demand in stratified coastal systems, such as Macquarie Harbour.

In this study, Macquarie Harbour was investigated as a model system to better understand the dynamics of microbial communities in semi-enclosed highly stratified aquatic environments. Water was sampled at two different depths, the surface and within the mixing zone, across the harbour to investigate how environmental parameters and the mixing zone affect the structure and composition of both the prokaryotic and eukaryotic communities using 16S and 18S rRNA gene-based high-throughput sequencing. This study (i) investigated the structure and distribution of both prokaryotic and eukaryotic microbial communities along the length of the Maquarie Harbour estuary, at the surface and within the mid-depth mixing zone; and (ii) explored how environmental gradients in salinity, temperature, chlorophyll-a and nutrients are related to microbial composition. We hypothesized that both prokaryotic and eukaryotic communities exhibit a common compositional variation as the concentration of the environmental parameters vary along the estuary as well as between the surface and within the mixing zone of the harbour. Also, we expect that the results would reveal patterns that could improve our knowledge of the ecology of the microbial communities.

## MATERIALS AND METHODS

### Location and sampling areas

Macquarie Harbour has a surface area of 276 km^2^, being approximately 33 km long and 9 km wide. Oceanic intrusion occurs over the sill at the mouth of the harbour through the Hells Gate strait (mean depth of 4 m and ~120m wide). The largest source of freshwater entering Macquarie Harbour is the Gordon River (GR), which flows into the south of the harbour, with mean monthly flows varying according to the season (< 100 m^3^ sec^-1^ in summer and early autumn, and 500 m^3^ sec^-1^ late autumn, winter, and spring) (Hartstein et al., 2019).

In November 2018, 46 water samples were collected using a 10 L Niskin bottle at three different locations spanning Macquarie Harbour from ocean to the Gordon River (GR) (Fig. 1). In Macquarie Harbour, water samples were taken from the surface (0-1 m; n= 16) and intermediate depths (4.5-11 m depth, n = 27), and from the surface only for the adjacent ocean (0-1 m; n = 3) and Gordon River (0-1 m; n = 4) samples. Samples were designated based on the water layer (S: surface, P: intermediate layer) and location (Macquarie Harbour: MH; Gordon River: GR; and ocean). Samples from within the harbour were also named according to the zone in which they were collected (A-C) (Fig.1). Samples were filtered for DNA extraction using a multichannel peristaltic pump. For each sample, two liters of water was filtered through 0.22 μM polyethersulfone Sterivex™ filter cartridges (Milipore, Darmstadt, Germany). Each sample was snap frozen and stored at −80°C until DNA extraction.

**Fig. 1.**
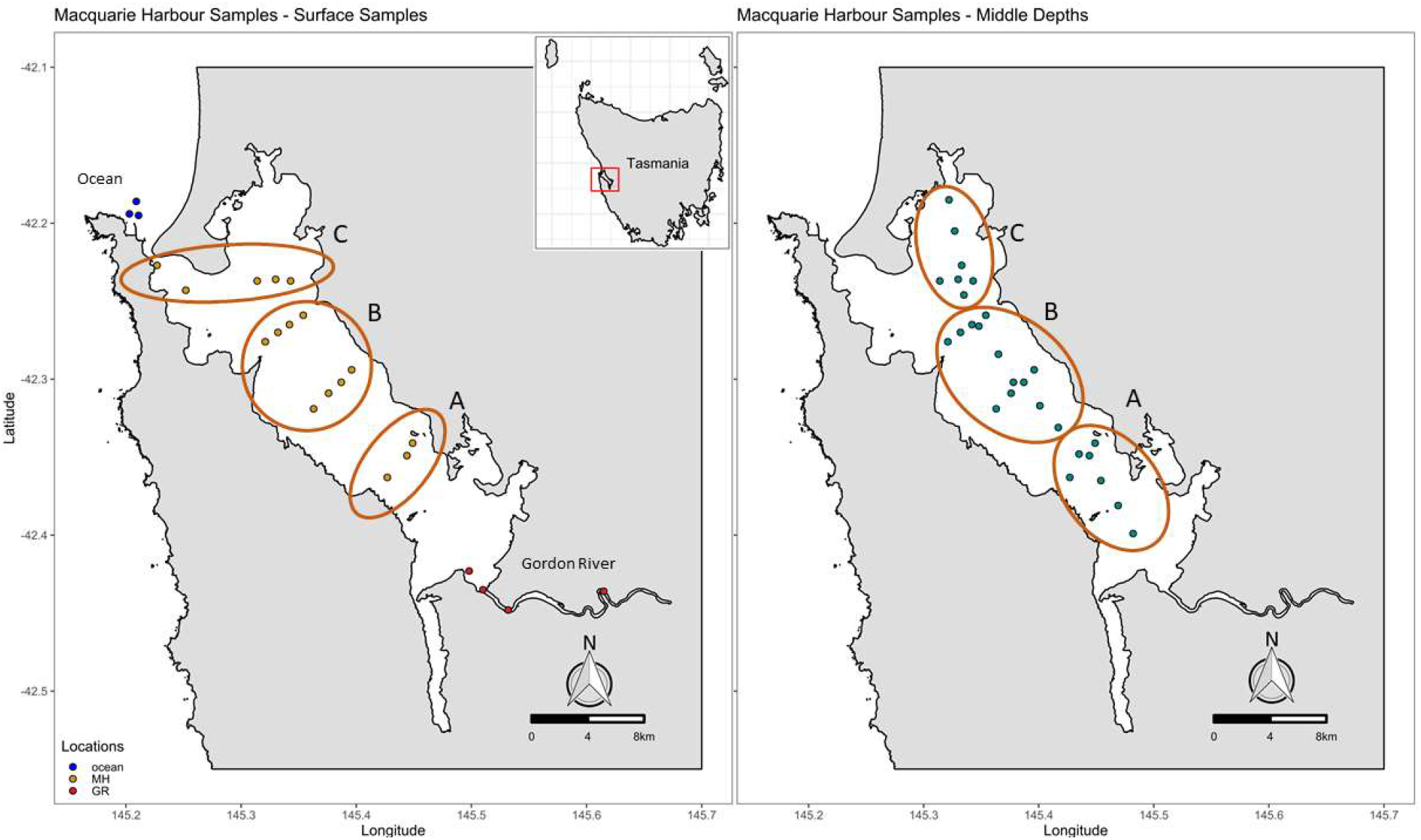
Study area showing the sample locations (dots). Surface waters were collected from the ocean (n = 3), Gordon River (n = 4), and Macquarie Harbour (n = 16). Intermediate waters were only sampled within the harbour (n = 27). The harbour was divided in three zones (A-C) to detect spatial and environmental gradients. S = Surface, P = middle waters.

### Environmental parameters

Environmental data included the sample location (latitude & longitude); temperature (°C), salinity, ammonium (NH_4_: μM), nitrate + nitrite (NO_x_: μM), nitrite (NO_2_: μM), phosphate (PO_4_: μM), silicate (SiO_4_: μM), and chlorophyll-a (Chl: μg/L). Physicochemical parameters were measured *in situ* at each sample location using a YSI 6600 V2 Multi Parameter Water Quality Sonde with a YSI 650 MDS logger. Water samples for dissolved inorganic nutrients and chlorophyll-a analyses were collected using a 10 L Niskin bottle sampler. Nutrient analyses were assayed using a Bran + Luebbe AA3 HR segmented flow analyzer following standard spectrophotometric methods (Grasshoff et al., 2009). Detection limits for NO_3_^-^/NO_2_^-^ were 0.015 μmol/L, for PO_4_^-3^ to 0.01 μmol/L, for NH_4_^+^ to 0.004 μmol/L, and for SiO_4_ to 0.01 μmol/L. For the chlorophyll-a (μg/L) determinations, water was extracted via gentle vacuum filtration (pressure drop <10 kPa) using 25-mm GF/F filters. Extracted chlorophyll-a samples were then measured on a Turner Trilogy laboratory fluorometer.

### DNA extraction and sequencing

DNA was extracted using a modified phenol:chloroform:isoamyl based DNA extraction protocol of the DNeasy PowerWater SterivexTM Kit (MoBio-Qiagen, vHilden, Germany) as described in Appleyard et al. (2013). Amplicon sequencing targeted the bacterial 16S rRNA gene V1-V3 region (27F–519R) (Lane et al., 1985; Lane, 1991), archaeal 16S rRNA gene V1-V3 region (A2F–519R) (Lane et al., 1985; DeLong, 1992) and eukaryotic 18S rRNA V4 region (TAReuk454FWD1-TAReuk-Rev3) (Stoeck et al., 2010). Libraries were separately generated and sequenced for each sample at the Ramaciotti Centre for Genomics (University of New South Wales, Sydney) on the MiSeq sequencing platform (Illumina Inc., USA) using 300 bp paired-end sequencing.

### Sequence analysis, OTU clustering and taxonomy assignment

Sequences were denoised to zero radius operational taxonomic units (zOTUS) as follows: Paired end reads were merged with FLASH2 (--min-overlap=30 --max-overlap=250) (Magoč and Salzberg, 2011). MOTHUR (v1.45.3) (Schloss et al., 2009) was used to remove sequences less than 400 bp, or containing N’s or homopolymer runs of > 8bp. The resultant sequences were then dereplicated (-fastx_uniques), sorted by read abundance (-sort_by_size) and denoised using the UNOISE3 algorithm (Edgar, 2016) in USEARCH (64-bit, v11.0.667) and default arguments. Merged reads were then mapped to the denoised zOTUs to produce a zOTU per sample abundance table (USEARCH -otutab). Taxonomic classification was performed with the SILVA 16S rRNA gene reference database version 138.1 for bacterial and archaeal communities (Yilmaz et al., 2014) and the SILVA 18S rRNA database version 128 and version 132 for eukaryotic community (Morien and Parfrey, 2018). Databases available at https://benjjneb.github.io/dada2/training.html. Using DADA2 (Callahan et al., 2016), taxonomic classification was performed with *assignTaxonomy* function (default arguments) which uses the naive Bayesian classifier method described in Wang et al. (2007). Raw sequence reads and metadata have been submitted to the NCBI Sequence Read Archive under bioproject PRJNA554530 (https://www.ncbi.nlm.nih.gov/bioproject/PRJNA554530/).

### Physicochemical analysis and alpha diversity Indices

To analyze differences in chemical environment between and among zones Principal Component Analysis (PCA) was performed on normalized data (standard deviation of one and a mean of zero). Prior to PCA, Spearman correlations were determined to search for highly correlated variables (i.e., r > 0.85, p-value ≤ 0.001). Permutational Multivariate Analysis of Variance (perMANOVA; (Anderson et al., 2011) and K-mean cluster analyses were used to examine the influences of water layers and zone on chemical measures of samples. Clustering validation was performed using the Elbow, Gap Statistics and Silhouette methods to ensure the optimal number of clusters (Thorndike, 1953; Rousseeuw, 1987; Tibshirani et al., 2001). Alpha diversity (Shannon diversity index and Inverse Simpson), species richness (Chao1) and evenness Pielou’s evenness index) were calculated. Prior to the analysis, sequence abundances for each sample was rarefied by adjusting differences in library size across samples (archaea = 40,000, bacteria = 59,116, eukaryotes = 95,000), by resampling with replacement (i.e., to equal sampling depth for each sample) using the *rarefy_even_depth* function of the phyloseq version 1.36.0 (McMurdie and Holmes, 2013). Statistical analyses in this step were conducted in R version 4.1.0 using the core distribution with the additional packages vegan version 2.5.7 (Oksanen et al., 2018) for PerMANOVA analysis, tidymodels version 0.1.3 (Kuhn and Wickham, 2020) for PCA and K-mean clustering, phyloseq version 1.36.0 (McMurdie and Holmes, 2013) and microbiome version 1.14.0 ((Leo Lahti and Shetty, 2017) packages to calculate diversity indices.

### Beta-Diversity and Drivers of Microbial Community

Multivariate analyses were used to examine the differences in microbial community compositions among locations and water layers, and to evaluate how environmental conditions affected the observed community variations. To focus on the core community and to reduce data complexity, zOTUs were filtered before analysis such that they represented a minimum of 0.1 % occurrence across all samples (Cao et al., 2021). Filtering resulted in core archaeal, bacterial, and eukaryotic communities of 521 zOTUs out of 1118 (2.02 % of total reads removed), 646 zOTUs out of 6015 (9.01 % of total reads removed), and 529 zOTUs out of 3116 (2.93 % of total reads removed), respectively. The abundance table was then centered log-ratio (CLR) transformed to address the negative correlation bias intrinsic to compositional data (Gloor et al., 2017). To visualize beta diversity among samples, distance-based RDA (db-RDA) plots were used on Aitchison distance (Euclidean distance on CLR-transformed data, Aitchison (1986) using the R package vegan (Oksanen et al., 2018). Both *envfit*, and *ordistep* functions from the vegan package, the latter with forward and backward stepwise selections, were used to examine significant relationships (p < 0.01 in at least one of the analyses) between microbial communities and environmental variables according to db-RDA. Next, PerMANOVA with 9999 permutations was used to test for significant differences in beta diversity among location and water layers (Gordon River surface, ocean surface, harbour surface, and harbour intermediate waters as factors). In the case of significant results (PerMANOVA = p < 0.01), pairwise PerMANOVA tests between groups with 9999 permutations were performed with Bonferroni significance adjustments to control for multiple pairwise comparisons. Permutation of dispersions was conducted to test if groups differed in their dispersion (Anderson, 2006) using the R package pairwiseAdonis version 0.4 (Martinez Arbizu, 2020). If significant differences were found, Tukey’s HSD with adjusted p-values were applied for multiple pairwise comparisons.

### Differential Abundance

To examine the major species that contributed to the differentiation between the sample groups revealed in the beta diversity analysis for each location, differential abundance analysis was performed using the R package Deseq2 version 1.32.0 (Love et al., 2014). The analysis was conducted on variance-stabilized data with a significance level set to 0.001 (McMurdie and Holmes, 2013).

### Co-occurrence networks and Partial least square regression

To investigate the interactions between members of microbial communities and environmental variables, weighted gene co-expression network analysis (WGCNA) was performed for each community separately using the WGCNA R package version 1.70.3 (Langfelder and Horvath, 2008). The analysis was based on previous studies investigating the relationship between microbial community structure and environmental data (Guidi et al., 2016; Horton et al., 2019). Briefly, the filtered and clr-transformed zOTU tables were used to generate a similarity matrix of nodes (i.e., zOTUs) based on pairwise Pearson correlations across samples. Next, an adjacency matrix (connection strength between two zOTUs) was calculated by raising the similarity matrix to a soft threshold power to ensure network scale-free topology (the nodes distribution and interaction) which is highly tolerant against errors in node connectivity (Albert et al., 2000; Zhang and Horvath, 2005). Network analyses were performed only for bacterial and eukaryotic communities (*p*; p = 14 for bacterial and p = 20 for eukaryotic communities), since the analysis failed to pick a soft-threshold of Pearson correlation coefficient to ensure a free-scale topology of the archaeal community network. Then, the adjacency matrix was used to create a topological overlap matrix (TOM), along with hierarchical clustering. TOM was applied to detect submodules (i.e., subnetworks) of highly co-correlated zOTUs within and outside the network. The first principal component of each submodule (i.e., eigenvalue) was Pearson correlated with each environmental variable (i.e., NOx, NO_2_, NH_4_, etc.). Subnetworks with the highest positive correlation (r > 0.75 and *p* value < 0.001) were then selected to explore the relationships among environmental factors and microbial community composition and structure.

In addition to network analysis, partial least square regression (PLS), using the PLS package version 2.8.0 (Wehrens and Mevik, 2007), was used to assess if a submodule structure (measured value) was capable of predicting the variability of an environmental parameter (response variable). PLS models were permutated 1000 times and Pearson correlations were performed between response variables and leave-one-out cross validation (LOOCV) predicted values. Both modelled and measured values were compared to analyze the explanatory power of the model. Further analysis was conducted if the regression found by the model was above the threshold of R^2^ = 0.5, while for R2 values below 0.5 the network was considered unable to predict a determined environmental variable. The importance of individual zOTUs to the PLS model was determined using the variable importance in projection (VIP) analysis (Chong and Jun, 2005). High VIP values suggest high importance in environmental factor prediction for that zOTU. In the analysis, a minimum correlation value was defined at different r values (between 0.2–0.5) for each network for both network construction and visualization. Individual correlations of each zOTU to a given variable and the number of co-correlations (i.e., node centrality) were plotted to visualize the relationship of the submodule structure to this environmental parameter.

## RESULTS

### Environmental characteristics

For surface waters most parameters displayed a spatial gradient from the river, through the estuary, to the ocean (Fig. 2). Chlorophyll-a (Chl) and ammonia-nitrogen (NH_4_) decreased from the Gordon River area (Chl: 26.77 ± 0.21 μg/L; NH_4_: 0.85 ± 0.01 μM) to the ocean (Chl: 12.57 ± 1.06 μg/L; NH_4_: 0.15 ± 0.02 μM). Phosphorous (PO_4_) and salinity followed an opposite gradient increasing from the river (PO_4_: 0.05 ± 0.005 μM; salinity: 0.22 ± 0.17) to the ocean (PO_4_: 0.29 ± 0.025 μM; salinity: 29.86 ± 1.47). Also, the samples from the ocean, water layers of the harbor and river showed significant variation in nutrients, temperature, salinity and chlorophyll (Fig. 2). Chl and NH_4_ were very low in the intermediate waters (4.5-11 m depth) (Chl: 0.62 ± 0.44 to 0.87+ 0.06 μg/L; NH_4_: 0.09 ± 0.09 to 0.12 ± 0.09 μM), which instead had the highest concentrations of nitrate/nitrite (NO_x_: 5.15 ± 0.56 to 5.85 ± 0.5 μM). Temperature and silicate (SiO_4_) were higher in the surface waters of the harbour (temperature: 17.78 ± 0.41 to 19.09 ± 0.58 °C; SiO_4_: 16.47 ± 0.15 μM to 15.90 ± 0.57 μM) compared with river (temperature: 12.32+ 0.43 °C, SiO_4_: 2.15 ± 0.97 μM), ocean (temperature: 15 ± 0.12 °C, SiO_4_: 5.37 ± 1.19 μM) and intermediate waters (temperature: ~15 ± 0.35 °C, SiO_4_: 15.26 ± 1.37 μM to 11.76 ± 0.53 μM). Salinity was higher in intermediate waters (16.64 ± 4.73 to 22.32+ 2.68) compared with the surface water of the harbour (6.55 ± 0.93 to 7.39 ± 2.85). Through PCA analysis on environmental data, salinity, NO_x_, NH_4_, chlorophyll and temperature were shown to be correlated with the spatial variability displayed in the ordination plot, in which the first two principal components explaining more than 75% of the variation of the original dataset (Fig.3). These two principal components clearly classified the samples in different groups according to location and depth (adjacent ocean, Gordon River, harbour surface and harbour intermediate waters). The result reinforces the spatial differences in environmental factors across the studied area.

**Fig 2.**
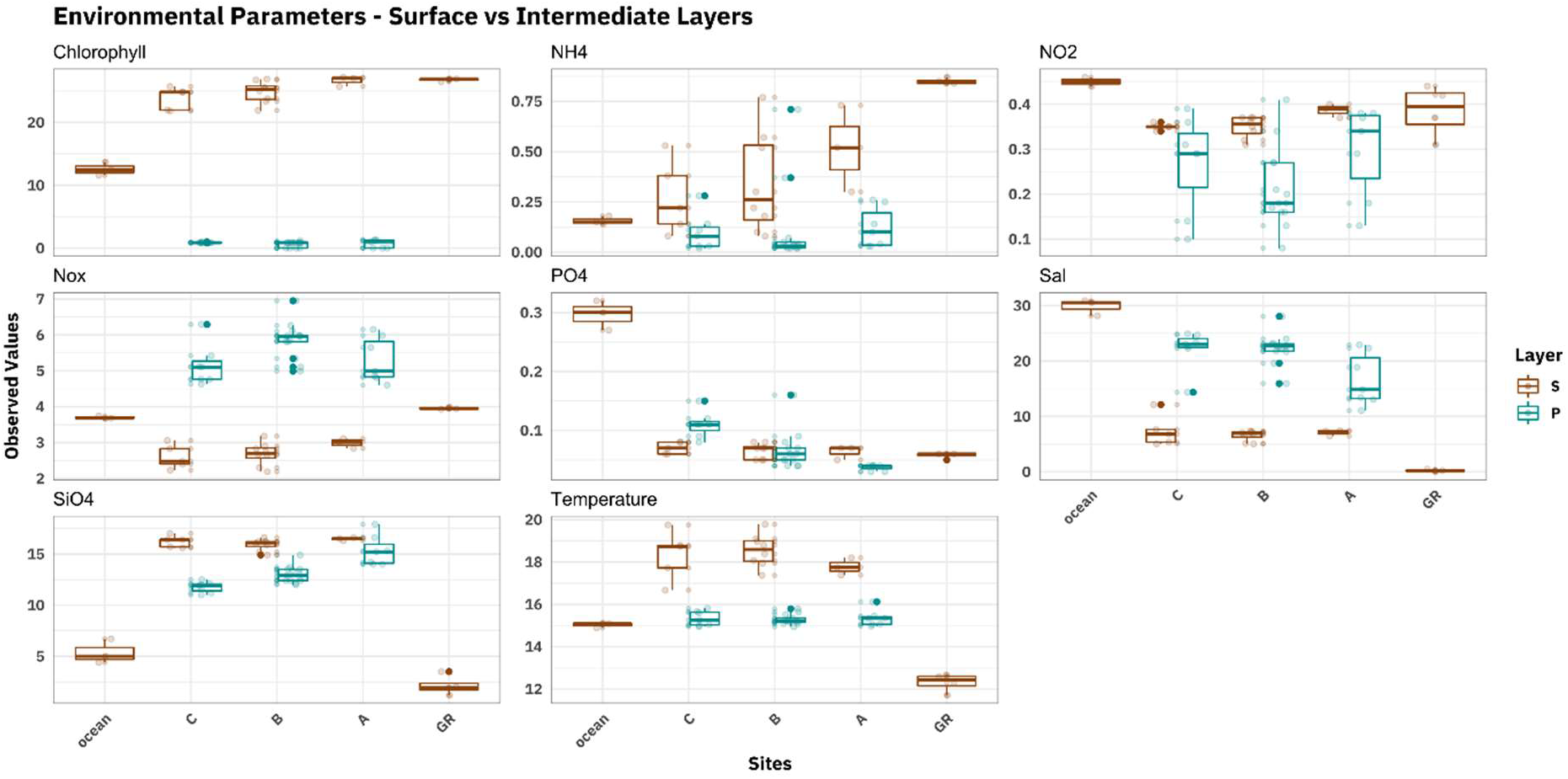
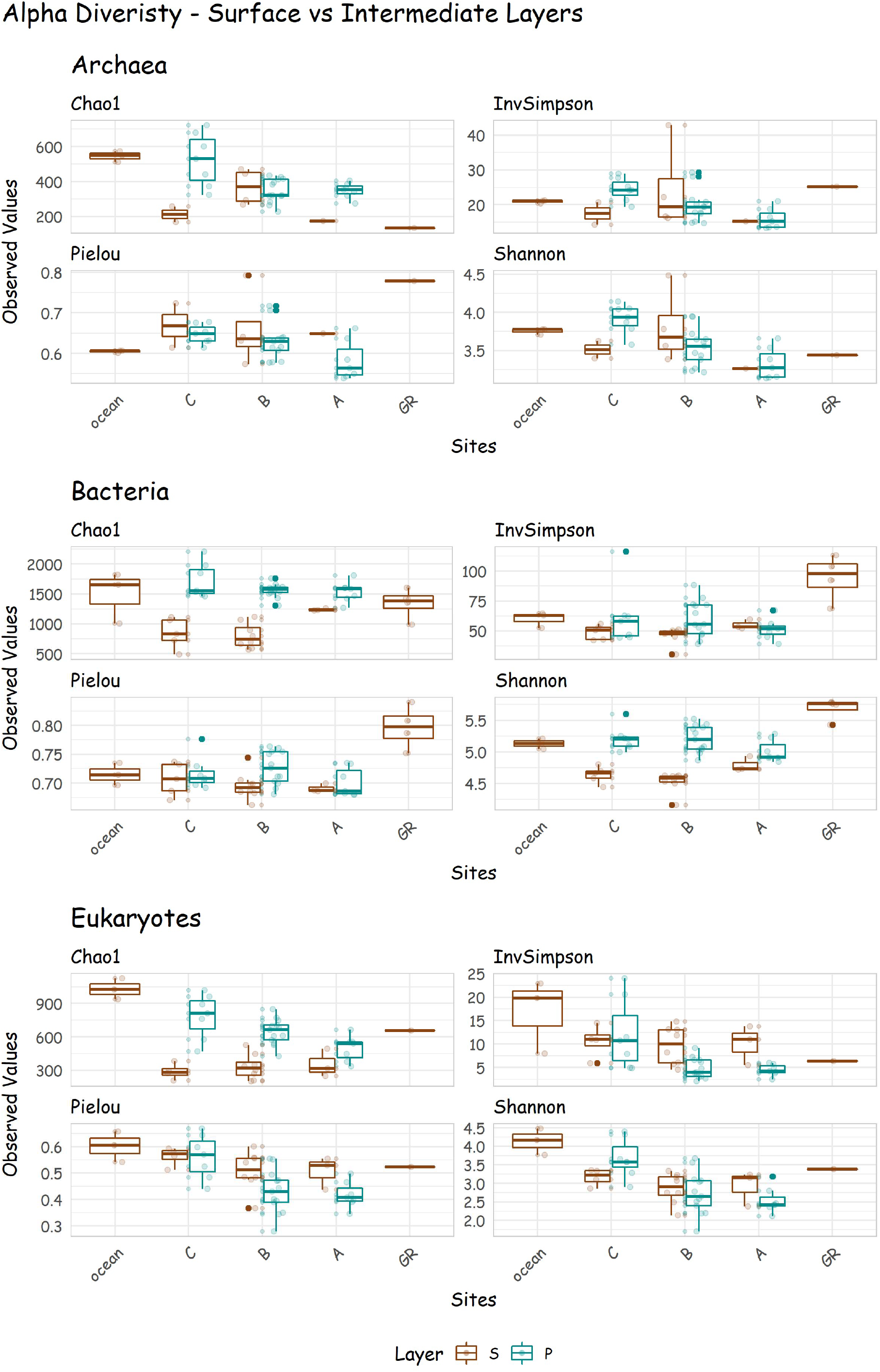
Box plots for environmental factors and diversity indices in different locations, and water layers. Temperature (°C), Sal: salinity (PSU), Chlorophyll (μg/L), and dissolved nutrients (μM). C, B, A refer to different zones of the harbour. GR refers to Gordon River. See Figure 1 for details.

**Fig 3.**
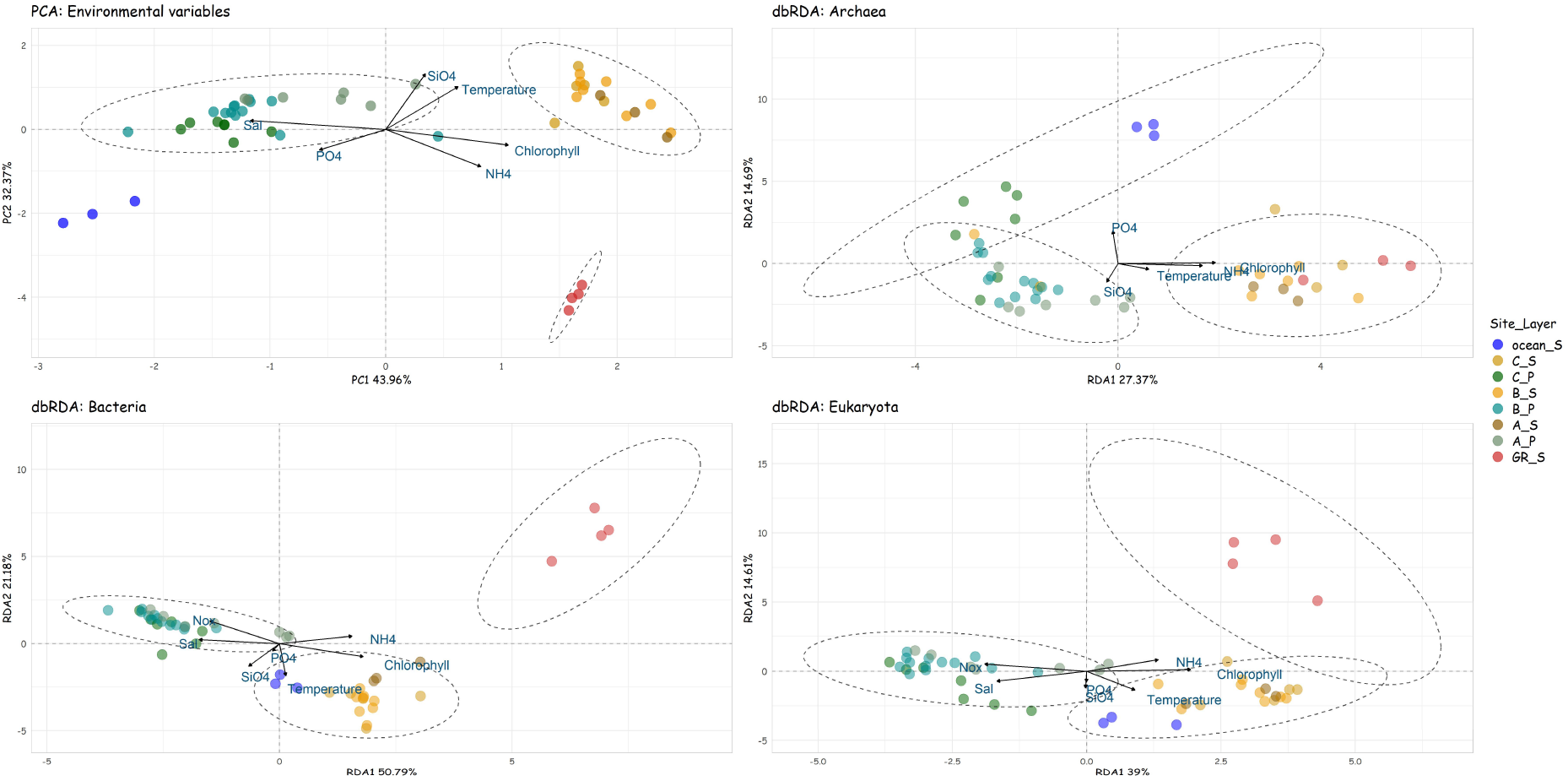
Principal Component Analysis (PCA) of environmental data and distance-based Redundancy Analysis (db-RDA) of the taxonomic differences of microbial communities. Dashed lines highlight the results of K-mean clustering analyses. Community analyses was based on the dissimilarities among microbial communities using the Aitchison distance. In the PCA analysis Salinity and NOx were combined due to high correlation. C, B, A refer to different zones of the harbour. GR refers to Gordon River. S refers to surface and P refers to intermediate water layer. See Figure 1 for details. Significant correlation between environmental variables and ordination were plotted as vectors (arrows) whose direction and length indicate the environmental variable gradient and the correlation, respectively.

### Richness and diversity

To assess the diversity, evenness, and richness of microbial communities through the different locations and water layers, Chao1, Inverse Simpson, Pielou and the Shannon diversity indexes were calculated (Fig. 2). The results indicate that microbial abundance and diversity changed along the environmental gradients of the harbour region. Archaeal estimated richness (Chao1) increased from the river to the ocean, whereas Pielou displayed an opposite pattern. For both bacterial and eukaryotic communities, the Chao1 was lower in the harbour surface waters. The bacterial diversity and evenness were higher in the Gordon river decreasing in the harbour water layers and ocean surface. The opposite trend was observed for the eukaryotic community, in which diversity and evenness increased from the river to the ocean.

### Microbial community composition and structure in different harbour locations

Community compositions in different locations and water layers were contrasted at the phylum, class and family levels (Fig. S1 - Fig. S6 Figs). The archaeal community included a total of 4 phyla, 7 classes and 5 families across all samples. *Crenarchaeota* was the most relatively abundant archaeal phylum in all locations and water layers. Also, *Thermoplasmatota* made a large contribution to the composition of ocean samples, while the phylum *Aenigmarchaeota* was present only in the samples from the Gordon River. At the family level, *Nitrosopumilaceae* was the most relatively abundant taxon. The samples from the Gordon River displayed a more even composition, with *Nitrosotaleaceae* and *Nitrososphaeraceae* also present. For the bacterial community, a total of 41 phyla, 85 classes and 268 families were found. *Proteobacteria* was the most relatively abundant phylum in all samples, followed by *Bacteroidota*, and *Actinobacteriota*. The phylum *Cyanobacteria* was relatively abundant in the surface waters of both the ocean and Macquarie Harbour. At the family level, *Flavobacteriaceae* (*Bacteroidia*) was relatively abundant in all samples except in the Gordon River, where *Comamonadaceae* (*Gammaproteobacteria*) was the most relatively abundant family. *Alphaproteobacteria* and *Gammaproteobacteria* were also abundant. SAR11 (*Alphaproteobacteria*) were relatively abundant in both ocean surface and intermediate layers of the harbour, while *Burkholderiaceae* and *Methylophilaceae*, (*Gammaproteobacteria*) were present in high relative abundance in river. In respect to the Eukaryotic community, 12 phyla, 36 classes and 261 families were detected. *Alveolata* (*Syndiniales*) and *Holozoa* (*Copepoda*) were the most relatively abundant groups in the samples from the ocean surface, while *Holozoa* (*Copepoda*) was the most relatively abundant at the surface of both Macquarie Harbour and Gordon River. On the other hand, the intermediate waters of the harbour were inhabited by a great number of Alveolata, encompassing the order *Syndiniales*. Members of *Stramenopiles* (or *Heterokonts*), including diatoms (*Ochrophyta*) and other forms of algae, were also found at all locations. This group was most relatively abundant in the ocean surface, but proportions decreased sharply in samples from the harbour, especially in the intermediate waters.

### Drivers of Microbial Communities

Beta-diversity analyses (db-RDA) revealed a separation of microbial communities by location and water layer and identified several environmental factors that were important in explaining community structure and distribution (Fig. 3). Archaeal communities were separated into two principal groups, one comprising the samples from the ocean and intermediate waters of the harbour and the other the samples from the river and harbour’s surface, which was confirmed by perMANOVA (R^2^ = 0.41, p < 0.01, betadisper < 0.01) and post hoc pairwise analysis (Table S2). Ordination analysis (db-RDA) explained 42% of the variation in archaeal community structure, which was significantly correlated to NH_4_, temperature and SiO_4_ (Table S3). The bacterial and eukaryotic communities showed similar results, with significant community separation between locations (ocean vs. harbour vs. river) and water layers (harbour surface vs. harbour intermediate waters), confirmed by perMANOVA (Bacteria: R^2^ = 0.75, p < 0.01; Eukaryotes: R^2^ = 0.59, p < 0.01) and pairwise post hoc analyses (Table S2). However, the non-homogeneous dispersion of the data could be influencing the results. In fact, we found different levels of dispersion by location and depth in the three microbial communities (*betadisper*, Archaea: *p* < 0.01, Bacteria: p = 0.01 and Eukaryotes: p = 0.01). While samples from the intermediate waters and river were grouped into two different clusters, samples from the ocean and the surface of the harbour were grouped together. Due to the small number of samples from the ocean (n = 3) and river (n = 4), post hoc analyses were unable to detect significant variation between the river and ocean samples. Db-RDA based on Aitchison distances analysis explained 42%, ~72% and 53% of the variation in the ordination of the archaeal, bacterial and eukaryotic communities, respectively. Silica (r^2^ env.fit = 0.37) was the most correlated environmental factor to this distinction between archaeal communities, temperature (r^2^ env.fit = 0.57) and SiO_4_ (r^2^ env.fit = 0.45) were the two top factors correlating for bacterial communities, while chlorophyll (r^2^ env.fit = 0.82) and salinity (r^2^ env.fit = 0.73) were the top two factors for the eukaryotic community (Table S3).

### Differential Abundance

Changes in community composition across sample depth and location were investigated using differential abundance analysis, with the main taxa driving changes similar to those observed through PCA and diversity analyses (Fig. S7-Fig. S9 Figs). Archaeal zOTUs assigned to *Nitrosopumilaceae* were detected in all locations by differential abundance analysis. *Nitrosotaleaceae* zOTUs were abundant in both surface of the harbour and river. *Nitrososphaeraceae* and *Thermoplasmatota* zOTUs were differentially abundant only in the river and ocean surfaces, respectively. *Syndiniales* (dinoflagellate), *Mediophyceae* (diatoms), *Mamiellales* (chlorophytes) and *Gymnodinium* (dinoflagellate) clades were differentially abundant in the ocean surface. The surface of the harbour was dominated by unique zOTUs from the family *Parvibaculaceae* of the class *Alphaproteobacteria*, along with a number of unique eukaryotic zOTUs belonging to the family *Copepoda*, followed by *Prostomatea*, and *Ochromonadales* but in smaller numbers. Intermediate water layer of the harbour and the Gordon River displayed different and unique bacterial and eukaryotic zOTUs, suggesting different communities inhabiting these environments (Fig. S8 and Fig. S9 Figs.). For instance, unique bacterial zOTUs from *Magnetospiraceae* (*Alphaproteobacteira*), *Pseudohongiellaceae* (*Gammaproteobacteira*), and Clade I (SAR11) as well as eukaryotic zOTUs from *Syndiniales, Chrysophyceae*, MAST-3A (i.e., marine *Stramenopiles*) among others were differentially abundant in the intermediate waters of the harbour. Additionally, bacterial members from *Comamonadaceae (Gammaproteobacteria), Burkholderiaceae (Gammaproteobacteria), Sporichthyaceae* (*Actinobacteria*) and *Chitinophagaceae* (*Bacteroidia*) along with eukaryotic zOTUs from *Chrysophyceae*, and *Peritrichia* were the most enriched families in the river surface.

### Network analysis

Having identified the environmental factors that are related to microbial community composition, further analyses were performed to attempt to identify specific microbial lineages affected by these variables (Fig. 3). From the network analysis, two bacterial (“blue” and “turquoise”, referred hereafter as bac1 and bac2, respectively) and two eukaryotic (“blue” and “brown”, referred hereafter as euk1 and euk2, respectively) submodules were identified for the measured dissolved nutrients (Fig Fig. S10, Fig. S11 and Fig. S12 and Table 2). Network analysis failed to pick a soft-threshold of Pearson correlation coefficient to ensure a free-scale topology of the archaeal community network. For this reason, only bacterial and eukaryotic network analyses were performed.

**Table 2.** Eigenvalue correlations (r > 0.75 and *p* value < 0.001) between microbial community submodules to nutrient variables, total zOTUs in each submodule and PLS model results (R^2^ > 0.5). Additional information can be found in Fig. S10, Fig. S11 and Fig. S12.

| Community | Nutrient | Submodule | Eigen correlations | Total zOTUs | PLSr | Measured x Predicted |
| --- | --- | --- | --- | --- | --- | --- |
| Bacterial | NO<sub>x</sub> | Blue/bac1 | R = 0.9 | 161 | R<sup>2</sup> = 0.88 | R = 0.95, p < 0.001 |
| Bacterial | NH<sub>4</sub> | Turquoise/bac2 | R = 0.79 | 248 | R<sup>2</sup> = 0.62 | R = 0.79, p < 0.001 |
| Eukaryotic | NO<sub>x</sub> | Blue/euk1 | R = 0.88 | 109 | R<sup>2</sup> = 0.78 | R = 0.89, p < 0.001 |
| Eukaryotic | PO<sub>4</sub> | Brown/euk2 | R = 0.89 | 45 | R<sup>2</sup> = 0.78 | R = 0.88, p < 0.001 |

Two submodules were found in bacterial and eukaryotic communities which were strongly related with NOx (Table 2; Fig. 4). The bac1 submodule comprised 161 zOTUs, most belonging to *Magnetospiraceae, Nitrospinaceae, Gimeciaceae* and *Bdellovibrionaceae* while the euk1 submodule was composed of 109 zOTUs most of which were related to *Syndiniales*. Both submodules are strongly correlated with NO_x_ (Fig. S13 and Fig. S14), suggesting that the zOTUs belonging to these two submodules are the most biologically related to that variable. Unclassified *Dehalococcoidia* (SAR202 clade), *Nitrospinaceae, Defluviicoccales, Syndiniales*, unclassified *Dinoflagellata*, and a novel *Cercozoa* Clade 2 were the most correlated bacterial and eukaryotic taxa to NO_x_, respectively (Table S4). Also, PLS analysis found that zOTUs from bac1 and euk1 submodules predicted 88% (Bacteria) and 78% (Eukaryotes) of NO_x_ variance (Table 2). According to VIP analysis, the zOTUs most strongly correlated with NOx concentration in the bacterial community were those related to *Dehalococcoidia* (SAR202 clade), *Magnetospiraceae* and *Bdellovibrionaceae* whereas in the eukaryotic communities were the ones encompassing *Syndiniales*, and *Dinoflagellata* (Table S5). The zOTUs with the highest connectivity belonged to the *Gracilibacteria* (*Patescibacteria*) within bacteria and to *Syndiniales* within the eukaryotes (Table S6).

**Fig 4.**
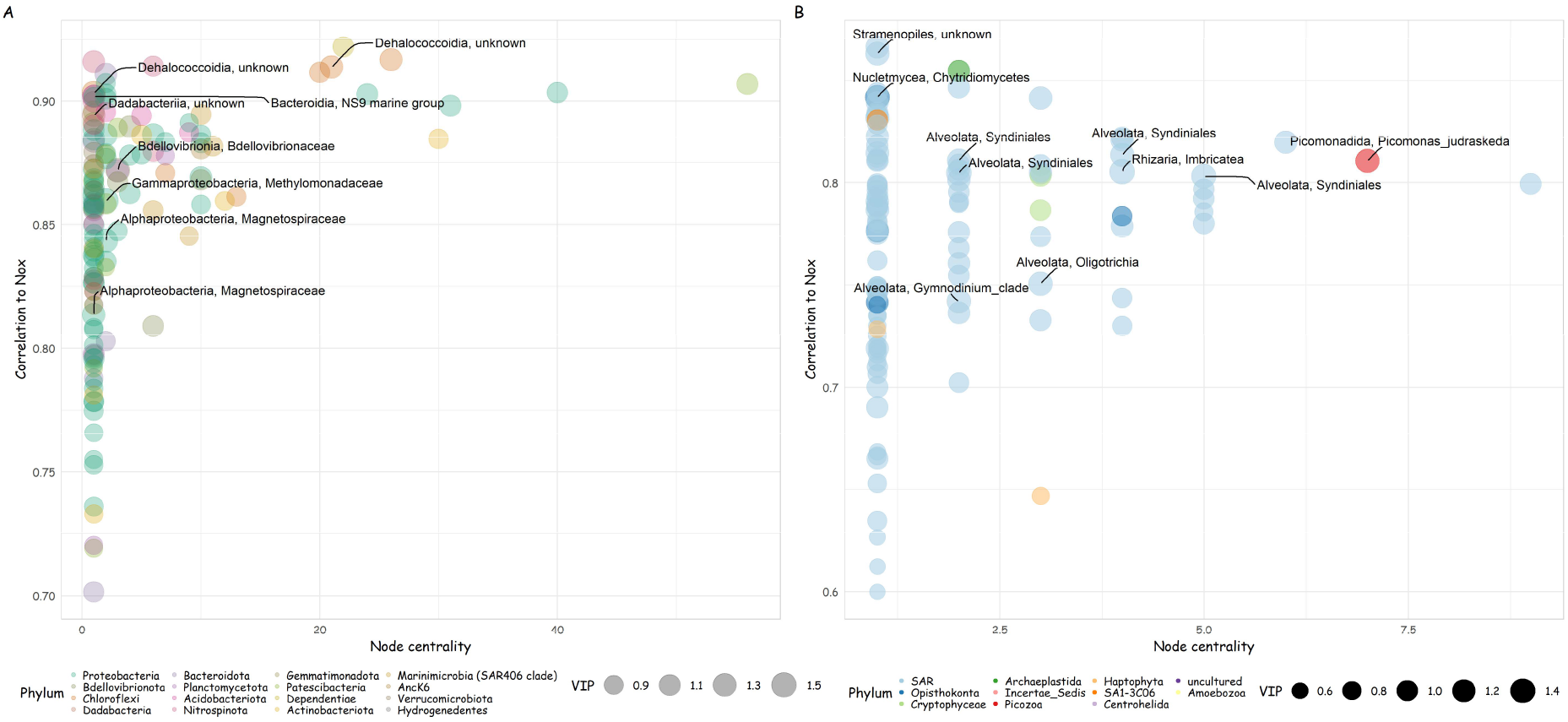
Bacterial (bac1) (A) and eukaryotic (euk1) (B) PLS results and submodules most correlated with NOx. The y-axis indicates zOTU correlation to NOx values, while the x-axis represents the number of co-correlations (node centrality). zOTUs are represented by points with color corresponds to the respective phylum. Point size is relative to VIP scores. The top zOTUs are identified with corresponding highest taxonomic classification possible.

The bac2 subnetwork was correlated to NH_4_ and had a prediction of 62% found by the PLS model (Table 2; Fig. S13). Among the 248 zOTUs grouped in this submodule, *Sporichthyaceae, Chitinophagaceae, Burkholderiaceae, Comamonadaceae*, and *Spirosomaceae* had the highest VIP scores and were frequently encountered. A *Bdellovibrionaceae* (OM27 clade) zOTU had the highest connectivity in this subnetwork linked to NH_4_ (Table S4 – Table S6 tables; Fig. 5 – A). The euk2 subnetwork had strong correlations to PO_4_. Only 45 zOTUs were grouped within this submodule which predicted 78% of the phosphate variation according to PLS results, albeit there was a low structural correlation to this parameter (Table 2; Fig. S14). The most correlated with PO_4_ and highest VIP scoring zOTUs encompassed *Syndiniales* and *Mediophyceae*, the latter had the highest number of co-correlations in the subnetwork (Table S4 – Table S6 tables; Fig. 5 – B).

**Fig 5.**
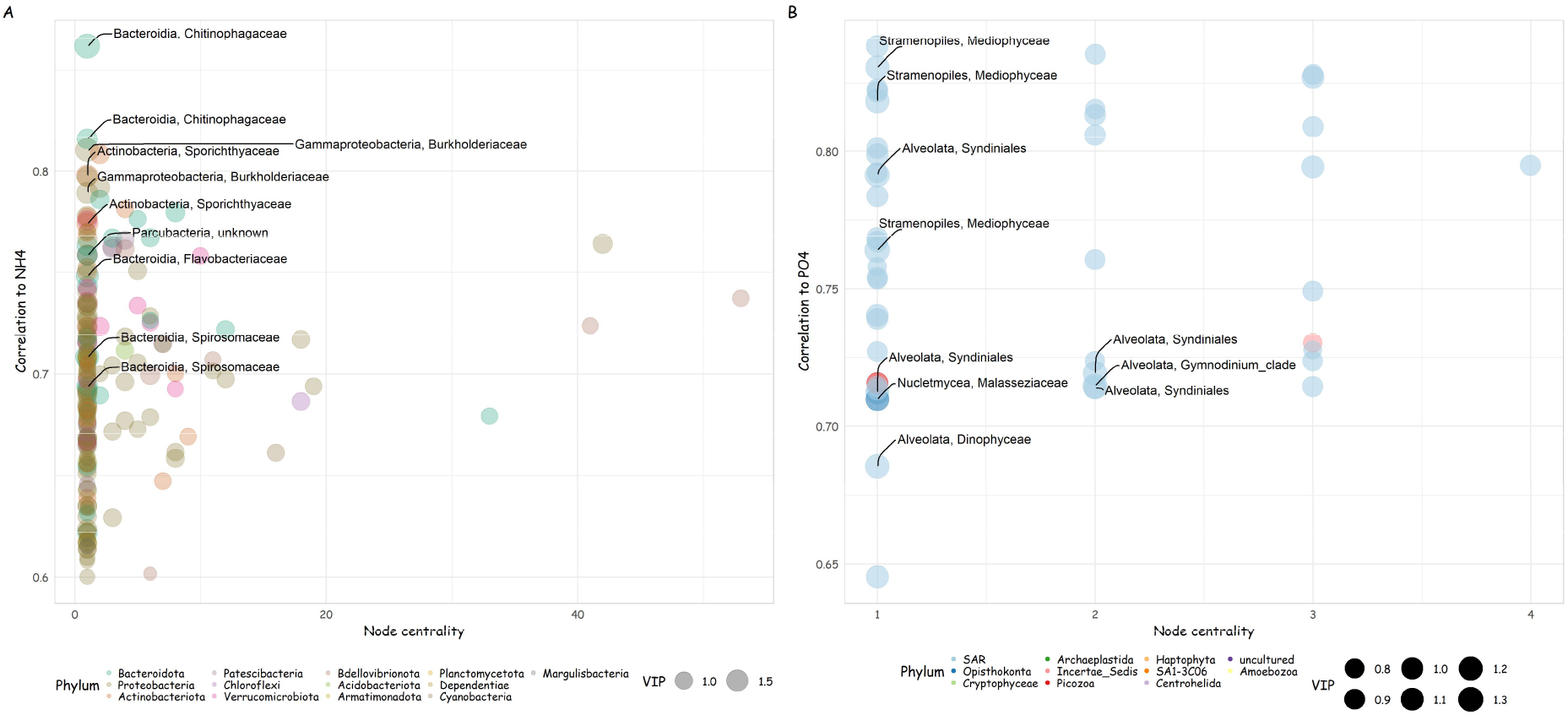
Bacterial (bac2) (A) and eukaryotic (euk2) (B) PLS results and submodules most correlated with NH_4_ and PO_4_, respectively. The y-axis indicates zOTU correlation to NH_4_ and PO_4_ values, while the x-axis represents the number of co-correlations (node centrality). zOTUs are represented by points with color corresponds to the respective phylum. Point size is relative to VIP scores. The top zOTUs are identified with corresponding highest taxonomic classification possible.

## DISCUSSION

The study region was characterized by broadscale physical and biochemical gradients in environmental parameters which drove microbial community structure. rRNA Gene amplicon sequencing, together with nutrients and other physiochemical data, allowed us to observe the structure and association of the microbial community in Macquarie Harbour. We found environmental parameters and microbial community composition varied according to location and depth. The results were not unexpected given the distinct longitudinal (river to ocean) and vertical (depth stratification) environmental gradients in the Harbour, supporting our hypothesis that that microbial community exhibits different compositions and distributions that largely follow physicochemical gradients along the estuary.

The changes in archaeal community compositions appeared to be linked to the environmental gradient along the estuary. Despite differences in the archaeal ordination showed in db-RDA, the ubiquitous ammonia oxidizing archaea (AOA) of the family *Nitrosopumilaceae* were the dominant group in the archaeal community at all locations. Within this family Candidatus *Nitrosopumilus* were the most relatively abundant in all locations, whereas *Nitrosarchaeum* appeared only in the harbour’s water layers being more relatively abundant at the surface. Both genera are known to be commonly found in estuarine and coastal waters (Mosier et al., 2012; Qin et al., 2017). *Thermoplasmatota* was characteristic of the oceanic community, and this group is globally distributed in ocean surface waters and, importantly, are likely photoheterotrophic, based on the presence of proteorhodopsin, enabling them to breakdown organic compounds using light as a supplementary energy source (Olson et al., 2018; Rinke et al., 2019). At the other end of the system *Nitrosotaleaceae*, *Nitrososphaeraceae*, and *Aenigmarchaeota* were characteristic of the river samples*. Nitrosotaleaceae* and *Nitrososphaeraceae* are ammonia oxidizers able to convert ammonia to nitrate (Alves et al., 2018; Zou et al., 2020), while *Aenigmarchaeota* are thought to be involved in carbon fixation, or rely on host-symbiont interactions for nutrient acquisition.(Castelle et al., 2015; Li et al., 2020). The archaeal community in the harbour surface water samples was more like the river samples, and this appears to be driven by the higher concentrations of NH_4_ emanating from the river along with the lacking of mixing with the subjacent water layer (Hitchcock and Mitrovic, 2015; Wymore et al., 2015). The archaeal community may be playing a major role in nutrient cycling in the Macquarie Harbour environment (Wuchter et al., 2006; Adam et al., 2017; Baker et al., 2020). Therefore, more studies are necessary to expand our knowledge on the ecology and physiological capabilities of this ecologically crucial group.

Bacterial and eukaryotic community compositions also varied across the major environments (river, ocean and harbour) and this appeared to be driven by changes in temperature, salinity, and dissolved nutrients. Although previous studies have described the transition in bacterial and eukaryotic communities in estuarine environments as a consequence of the mixing between river and ocean (Herlemann et al., 2011; Li et al., 2018; Anas et al., 2021) others have shown that coastal environments are inhabited by a distinct community structure adapted to the specific environmental conditions (Storesund et al., 2017; Ghosh and Bhadury, 2019; Gong et al., 2020). This study shows that in Macquarie Harbour the distributions of both bacterial and eukaryotic communities are influenced by physicochemical and biological gradients where microbial communities are adapted to specific environmental conditions.

The Gordon River samples were characterized by a distinct bacterial and eukaryotic communities. The bacterial community featured the taxa *Comamonadaceae*, *Burkholderiaceae* (both from order *Burkholderiales*), *Sporichthyaceae* (*Actinobacteria*) and *Chitinophagaceae* (*Bacteroidia*). These taxa were also the most strongly related to NH_4_, as evidenced through both network analysis and PLS model. These bacterial groups that correlate to NH_4_ concentrations include putative copiotrophs which are common to many environments, such as soils, rivers and estuarine waters (Carney et al., 2015; Aylagas et al., 2017; Griffin et al., 2017). Additionally, copiotrophs are also known to become dominant in eutrophic waters (Chenard et al., 2019) or in the surface of coastal ocean regions with high rates of organic matter degradation (Golebiewski et al., 2017; He et al., 2017; Sörenson et al., 2020). Network analysis revealed *Bdellovibrionaceae* as the node with the highest node centrality in the bac2 subnetwork. *Bdellovibrionaceae* are defined as obligate predators, and as such, their position in the subnetwork is consistent with the group potentially controlling the bacterial community. This bacterial prey-predator relationship has also been reported in others microbial networks from lacustrine systems (Ezzedine et al., 2020) and paddy soils (Qian et al., 2020). Regarding the eukaryotic community, several taxonomic groups were differentially abundant among riverine samples. Chrysophyceae and Peritrichia were the most enriched taxa at this site. Generally, chrysophytes are considered ubiquitous with high plasticity in energy source use (Choi et al., 2016; Henson et al., 2018) and can rely on bacterial grazing, especially in freshwater (Rottberger et al., 2013). Moreover, the presence of these two eukaryotic groups could be related to the high amount of dissolved and particulate organic matter in the Gordon River which is tannin-rich (Terry, 2001; Dias et al., 2008; Rottberger et al., 2013; Hartstein et al., 2016). Also, Calanoida (Copepoda) made up a substantial proportion of the eukaryotic community in the river waters, most likely as primary consumers in response to the high levels of chlorophyll present (Turner, 2004; Kwok et al., 2015). Although inhabiting distinct bacterial and eukaryotic communities, it is clear that the Gordon River waters have a strong influence on environmental parameters in the Harbour and subsequently likely plays a large role in driving microbial interactions therein.

The bacterial and eukaryotic communities in the subsurface waters of the Macquarie Harbour were highly correlated to NO_x_ concentrations. Among bacterial zOTUs, PLS model and network analyses revealed that *Magnetospiraceae, Dehalococcoidia*, and *Nitrospinaceae* are correlated to nitrate concentration. *Magnetospiraceae* have been associated with nitrate reduction (Li et al., 2012; Bazylinski et al., 2013). *Dehalococcoidia* (SAR202 clade) are known to be key players in carbon and sulfur cycling in aphotic zones, contributing to the degradation of recalcitrant organic compounds forming syntrophic networks with organic matter degraders (Landry et al., 2017; Yang et al., 2020; Cabello-Yeves et al., 2021). Nitrite oxidizers, *Nitrospinaceae*, are one of the important players in the nitrogen cycle, as observed at the redoxcline and in oxygen minimum zones of the Black Sea and the eastern tropical North Pacific Ocean, respectively (Beman et al., 2013; Cabello-Yeves et al., 2020), where metabolic coupling with NH_4_-oxidizing Archaea can also occur (Mincer et al., 2007; Santoro et al., 2010). The association of these zOTUs with high nitrate concentration, along with low DO in the intermediate waters, indicates that these microorganisms are likely key players in nutrient cycling and further raises the possibility that they may play a role in the oxygen demand inside the harbour.

Within the Eukaryotes, *Syndiniales* was abundant in the harbour samples and highly related to NO_x_ concentrations. This group of alveolates also often dominates the pelagic zone (Clarke et al., 2019) and is well known for its parasitic lifestyle killing protistan and metazoan hosts, including dinoflagellates, ciliates, and copepods (Chambouvet et al., 2008; Zamora-Terol et al., 2020). The presence of zOTUS associated with dinoflagellates such as *Gymnodinium* and the observed dominance of copepods suggest that the presence of *Syndiniales* is consistent with its parasitic lifestyle reported in the marine environment (De Vargas et al., 2015) in locations such as the Baltic Sea (Zamora-Terol et al., 2020) and other stratified systems (Parris et al., 2014; Jing et al., 2015; Torres-Beltrán et al., 2018). We argue that the high abundance of *Syndiniales* in intermediate waters could be related to the favorable environmental conditions (e.g., high nutrient concentration and abundance of hosts) that allow this group to thrive when it enters via oceanic waters into the harbour. Parasitic infection, unlike predation, can affect the fate of nutrients in the food web by redirecting host biomass (source of organic matter) to the microbial loop rather than to higher trophic levels as would occur via predation (Yih and Coats, 2000; Torres-Beltrán et al., 2018). This pathway would also likely fuel the consumption of oxygen by the bacterial community. Therefore, these marine alveolates could be playing major roles in the structure and connectivity of food webs as well as the biogeochemical cycle in the water column of the harbour (Arístegui et al., 2009; Fuhrman et al., 2015; Anderson and Harvey, 2020) and thus should be considered in nutrient flux models.

This study also attempted to help better understand the drivers of the low DO commonly detected below the surface in the water column of the Macquarie Harbour, a tannin rich, stratified and semi-enclosed water system. The findings highlight that the harbour is inhabited by a diverse microbial community composed of oceanic and riverine taxa mixed along an environmental gradient, as well as taxa unique to the Harbour. Network and machine learning analyses revealed subcommunities occurring in the water column, particularly in the intermediate waters and in the Gordon River. Bacterial and eukaryotic communities inhabiting intermediate waters were related to NO_x_ concentration (submodules blue, bac1 and euk1) and the bacterial community from the river were associated to ammonia concentration (bac2 subnetwork). The differences detected in environmental variables (i.e., spatial gradient) and microbial distribution suggest that the harbour’s microbial populations may be stimulated by the harbour’s primary freshwater source, the Gordon River, which is known to transport both labile and recalcitrant organic matter along with the corresponding microbial community into Macquarie Harbour (Carpenter et al., 1991; Teasdale et al., 2003; Da Silva et al., 2021). Moreover, tannin-colored waters affect light penetration impacting phytoplankton biomass and phototrophic activity (Harding Jr et al., 1986; Edgar et al., 1999; Bledsoe et al., 2004; Cloern et al., 2014). In the presence of riverine inflows and allochthonous organic matter, both nitrogen and phosphorous (P) can act as limiting factors, changing the relationships between heterotrophic bacteria and phototrophic phytoplankton in which the former can outcompete the latter for the same limited nutrients, affecting the structure of the microbial community in the harbour (Currie and Kalff, 1984; Thingstad and Pengerud, 1985; Cotner et al., 1997; Grover, 2000; Mindl et al., 2005; Hitchcock and Mitrovic, 2013; Sörenson et al., 2020). However, more research regarding the relationship between nutrient loading and bacteria-phytoplankton interaction is necessary to confirm this assumption.

In addition to the harbour’s natural characteristics, the microbial communities inhabiting the surface and subjacent water layer likely contribute to the nutrient cycling and biogeochemical processes in this estuarine mixing-zone. The estuary may be inhabited by microorganisms adapted to the rapid oxidation of organic matter when oxygen becomes available (Lauro et al., 2009; Martens-Habbena et al., 2009; Jiao et al., 2014; Wild-Allen et al., 2020). Additionally, differential abundance and network analyses also provided important insights on the interactions occurring within the harbour. For instance, some important zOTUS in the subnetworks and locations were assigned to taxa commonly suggested as predators or parasites which have been associated to changes in carbon and nutrients fluxes in aquatic food webs (Worden et al., 2015). Then these zOTUS highly correlated to NO_x_ and ammonia could be playing important role in the nutrient cycle in the harbour and as a result may have potential as indicators of environmental health (Paerl et al., 2003; Pawlowski et al., 2016). Therefore, these observed associations and taxonomic groups provide baseline information that can be incorporated into future studies on biogeochemical cycle and oxygen demand in Macquarie Harbour and similar regions. This study represents a first step in offering more information on the biological complexity for better understand ecological processes in stratified coastal or estuarine areas.

