## supplementary figures for "The importance of environmental parameters and mixing zone in shaping estuarine microbial communities along a freshwater-marine gradient"

### SUPPLEMENTARY MATERIAL - FIGURES

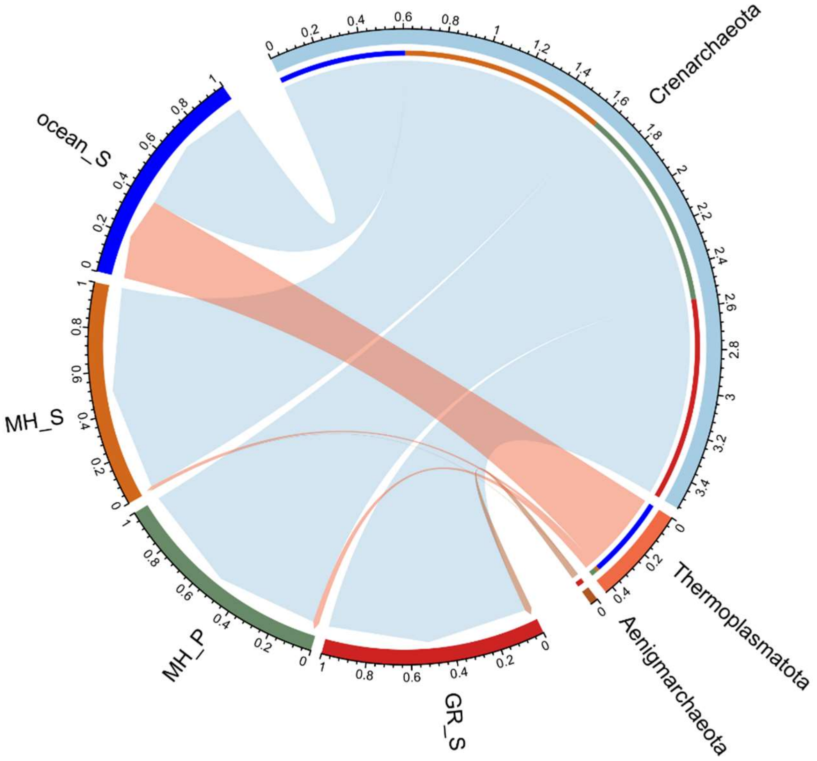


Fig. S1 - Archaeal composition at phylum level. The circus plot shows the community structure of all water samples at the phylum level. Data was filtered before the analysis.

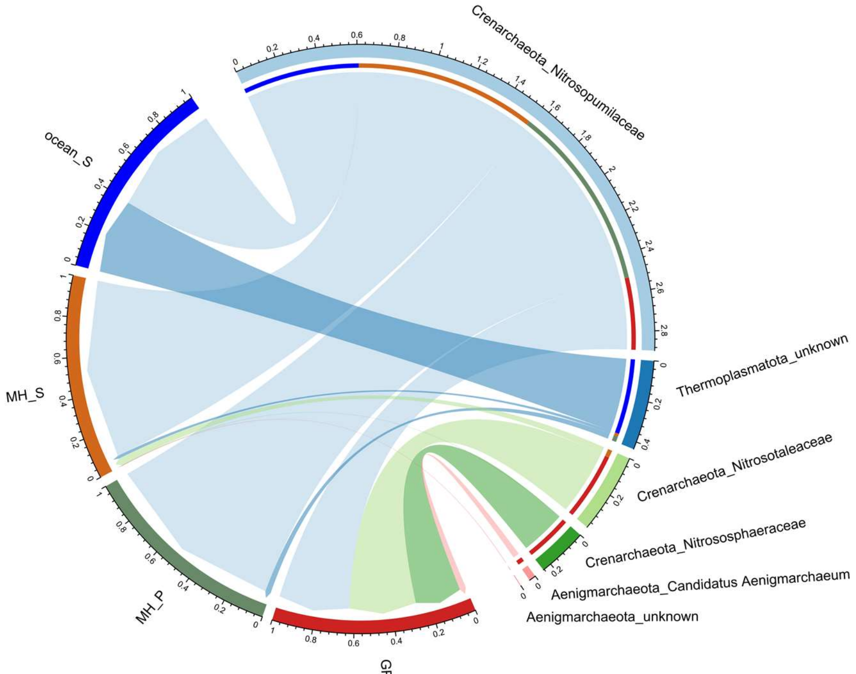


Fig. S2 - Archaeal composition at family level. The circus plot showing the community structure of all water samples at the phylum level. Data was filtered before the analysis.

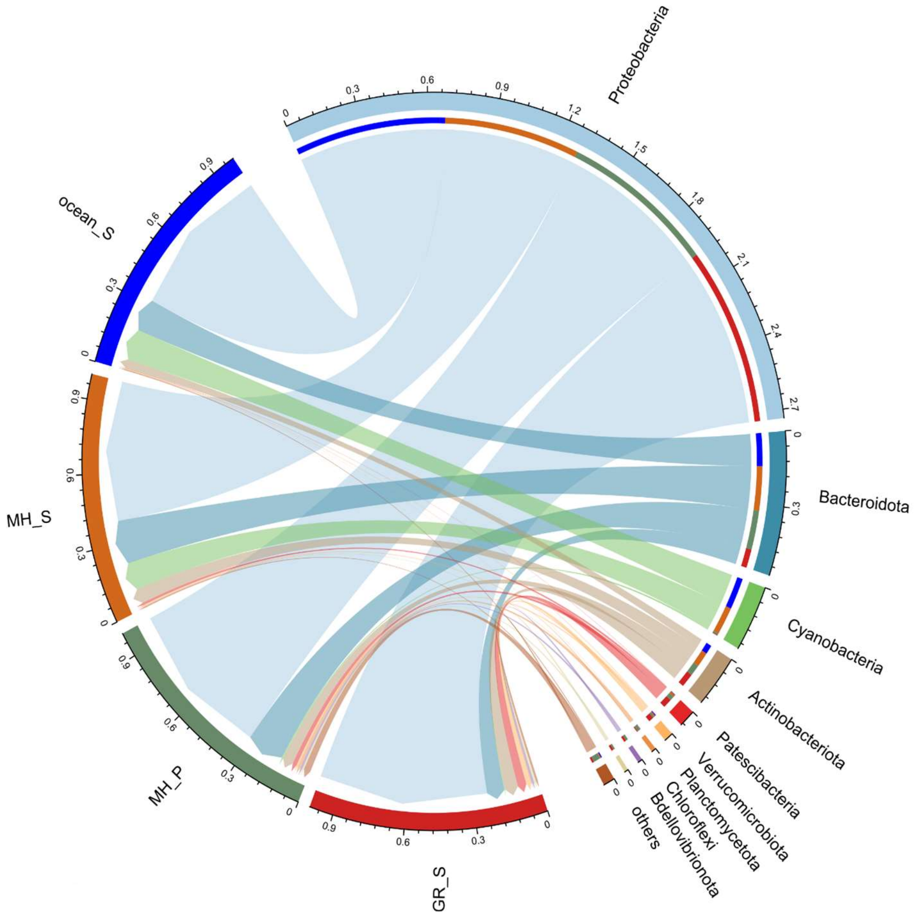


Fig. S3 - Bacterial composition at phylum level. The circus plot shows the community structure of all water samples at the phylum level. Data was filtered before the analysis. For visualization purpose, only the more abundant taxa were displayed. Less abundant taxa were grouped as “others”.

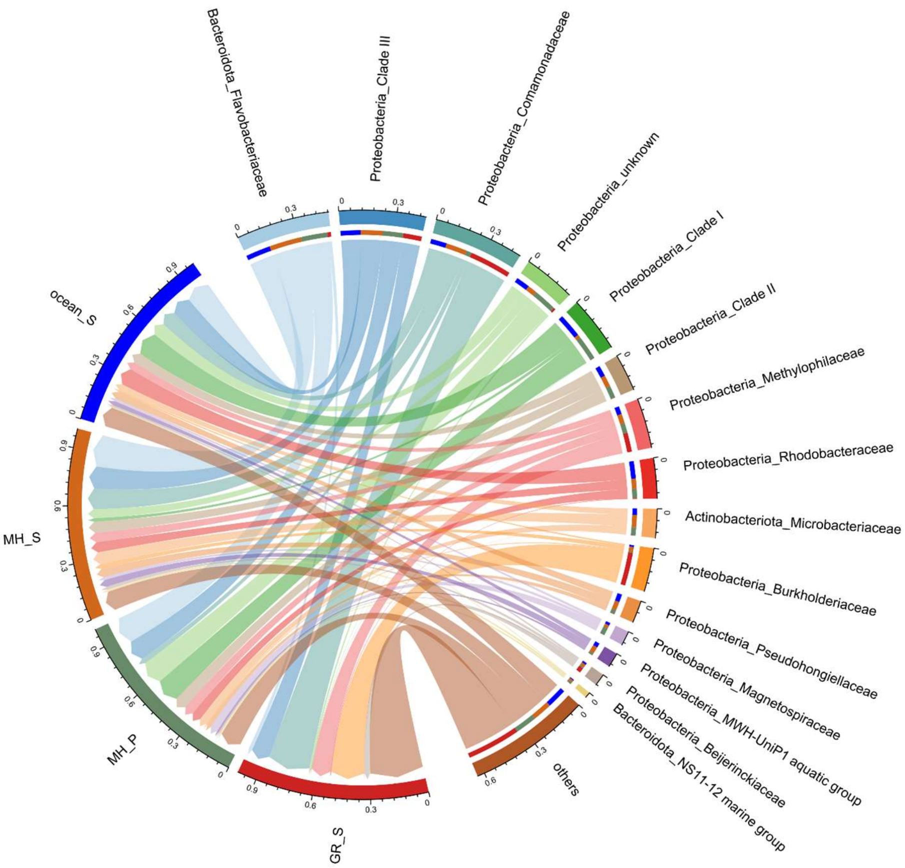


Fig. S4 - Bacterial composition at family level. The circus plot shows the community structure of all water samples at the phylum level. Data was filtered before the analysis. For visualization purpose, only the more abundant taxa were displayed. Less abundant taxa were grouped as “others”.

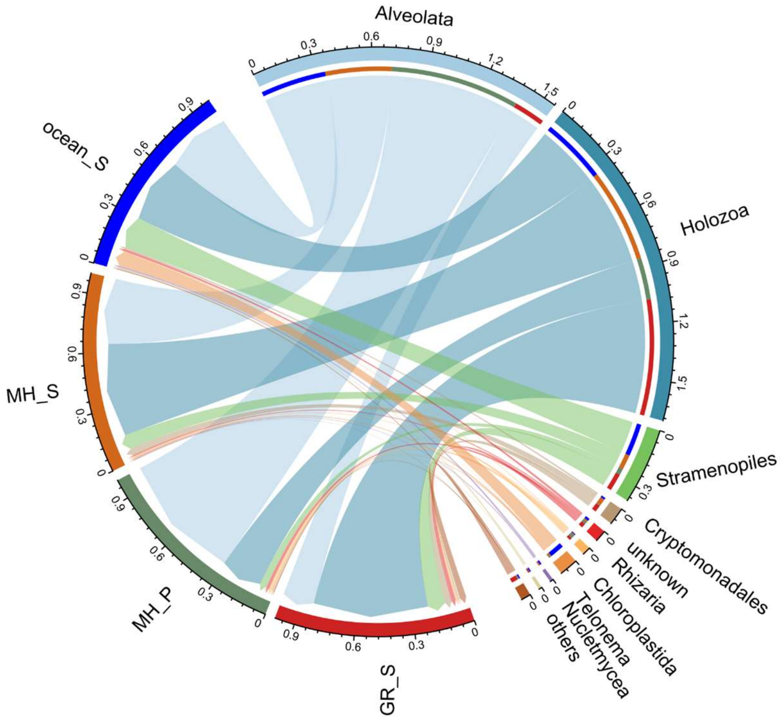


Fig. S5 - Eukaryotic composition at class level. The circus plot shows the community structure of all water samples at the phylum level. Data was filtered before the analysis. For visualization purpose, only the more abundant taxa were displayed. Less abundant taxa were grouped as “others”.

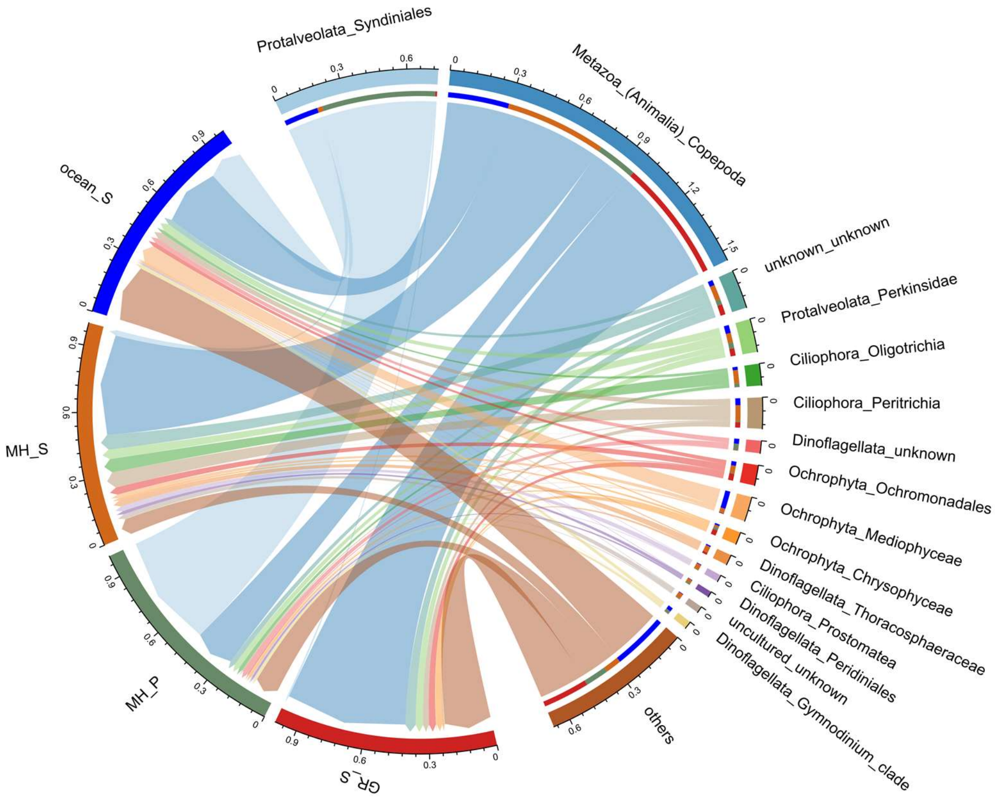


Fig. S6 - Eukaryotic composition at family level. The circus plot shows the community structure of all water samples at the order level. Data was filtered before the analysis. For visualization purpose, only the more abundant taxa were displayed. Less abundant taxa were grouped as "others".

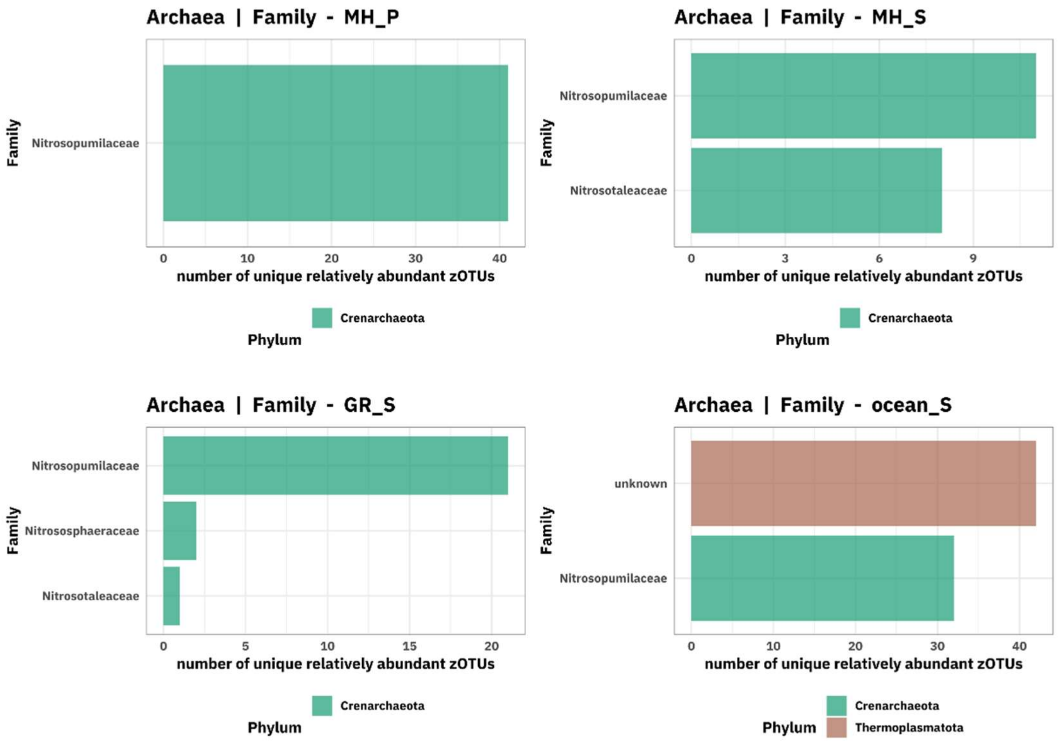


Fig. S7 - Differentially abundant archaeal zOTUs at family level. Only unique zOTUs for each site with a significance level of 0.001 and with a log2 fold-change more than |2| for each site are shown. Each bar indicates the total number of zOTUs belonging to each taxonomic level that was uniquely enriched in the respective site after the comparison with all the other sites. Different color represents different phyla.

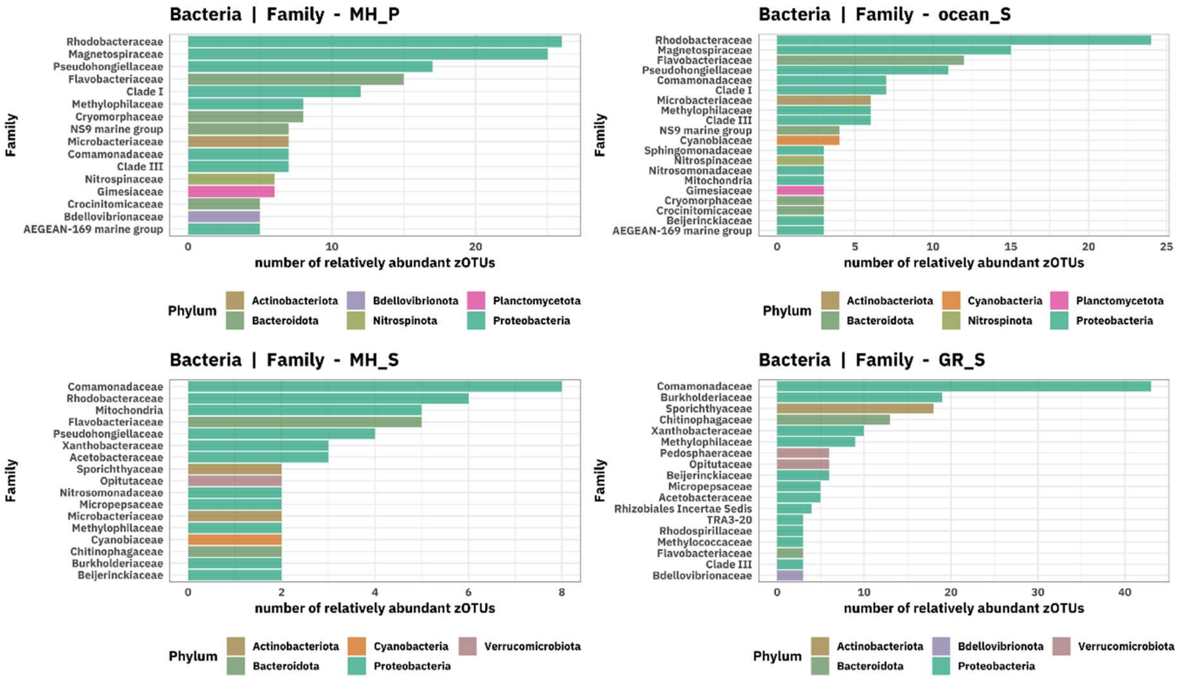


Fig. S8 - Differentially abundant bacterial zOTUs at family level. Only unique zOTUs for each site with a significance level of 0.001 and with a log2 fold-change more than |2| for each site are shown. Each bar indicates the total number of zOTUs belonging to each taxonomic level that was uniquely enriched in the respective site after the comparison with all the other sites. Different color represents different phyla.

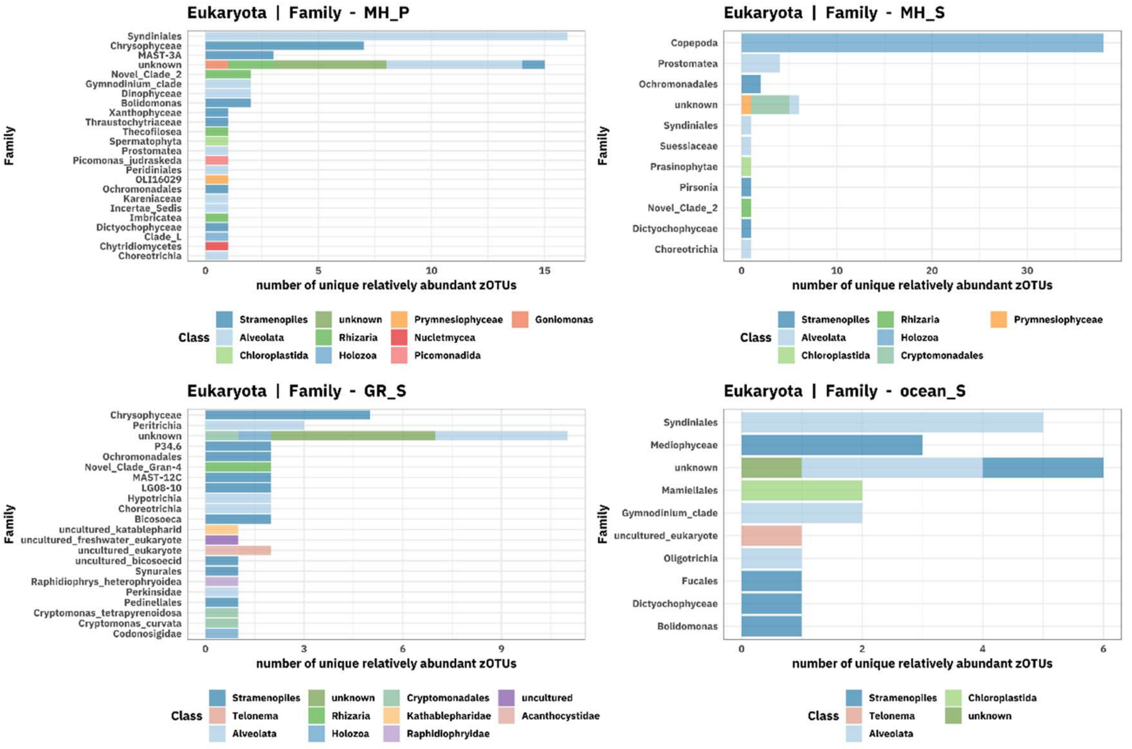


Fig. S9 - Differentially abundant eukaryotic zOTUs at family level. Only unique zOTUs for each site with a significance level of 0.001 and with a log2 fold-change more than |2| for each site are shown. Each bar indicates the total number of zOTUs belonging to each taxonomic level that was uniquely enriched in the respective site after the comparison with all the other sites. Different color represents different class.

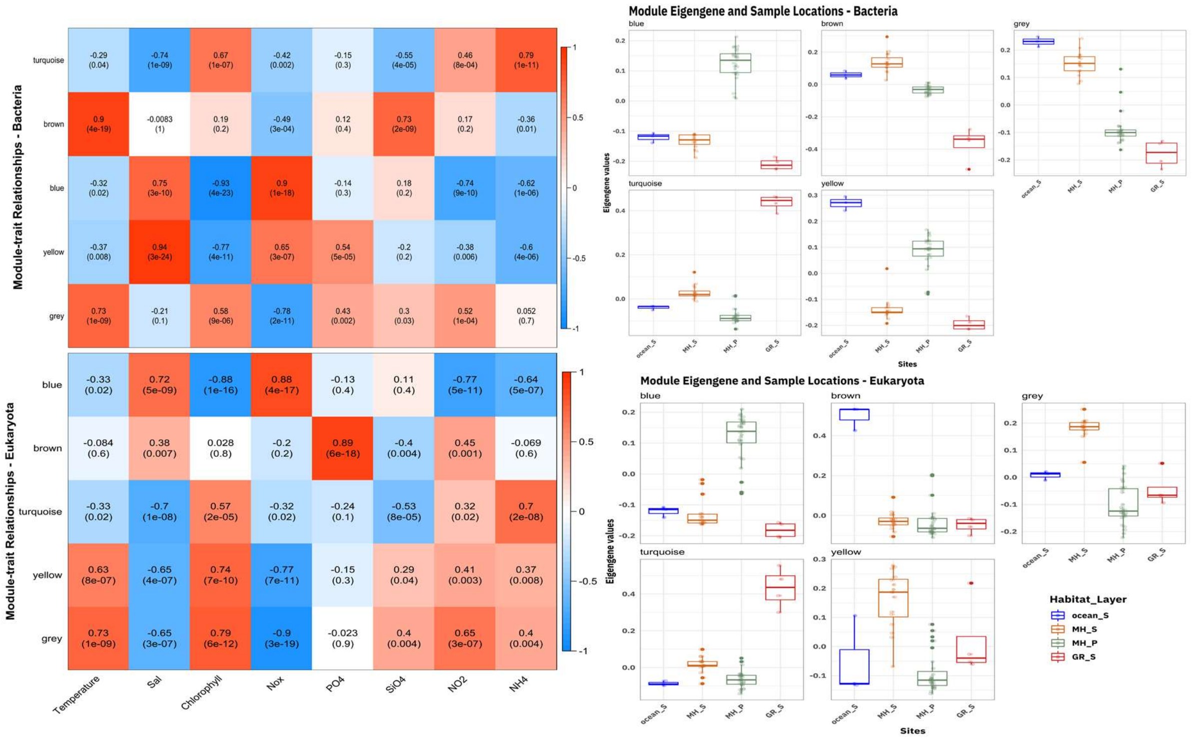


Fig. S10 - Relationships of consensus module eigengenes and traits (environmental variable). Each row in the table corresponds to a consensus module (of zOTUs), and each column is an environmental variable. Numbers in the table report the correlations of the corresponding module eigengenes and traits, with a p-value printed below in parentheses. The table is color coded by correlation according to the legend. From the plot above, it is possible to identify the modules that are significantly correlated to each environmental variable (pvalues in the parentheses). Also, boxplot representations of module eigengene values which can separate samples based on site and water layer. Module eigengene is a single representative expression profile for each sample, based on the principal component of that module. The boxplots display the blue eigengene values separating the intermediate water samples while the turquoise eigengene values could distinct the river samples from the others for both bacterial and eukaryotic communities. The samples from the ocean were grouped in yellow and brown modules eigengenes for bacterial and eukaryotic communities, respectively. The grey module is where all zOTUs that do not fall into other modules.

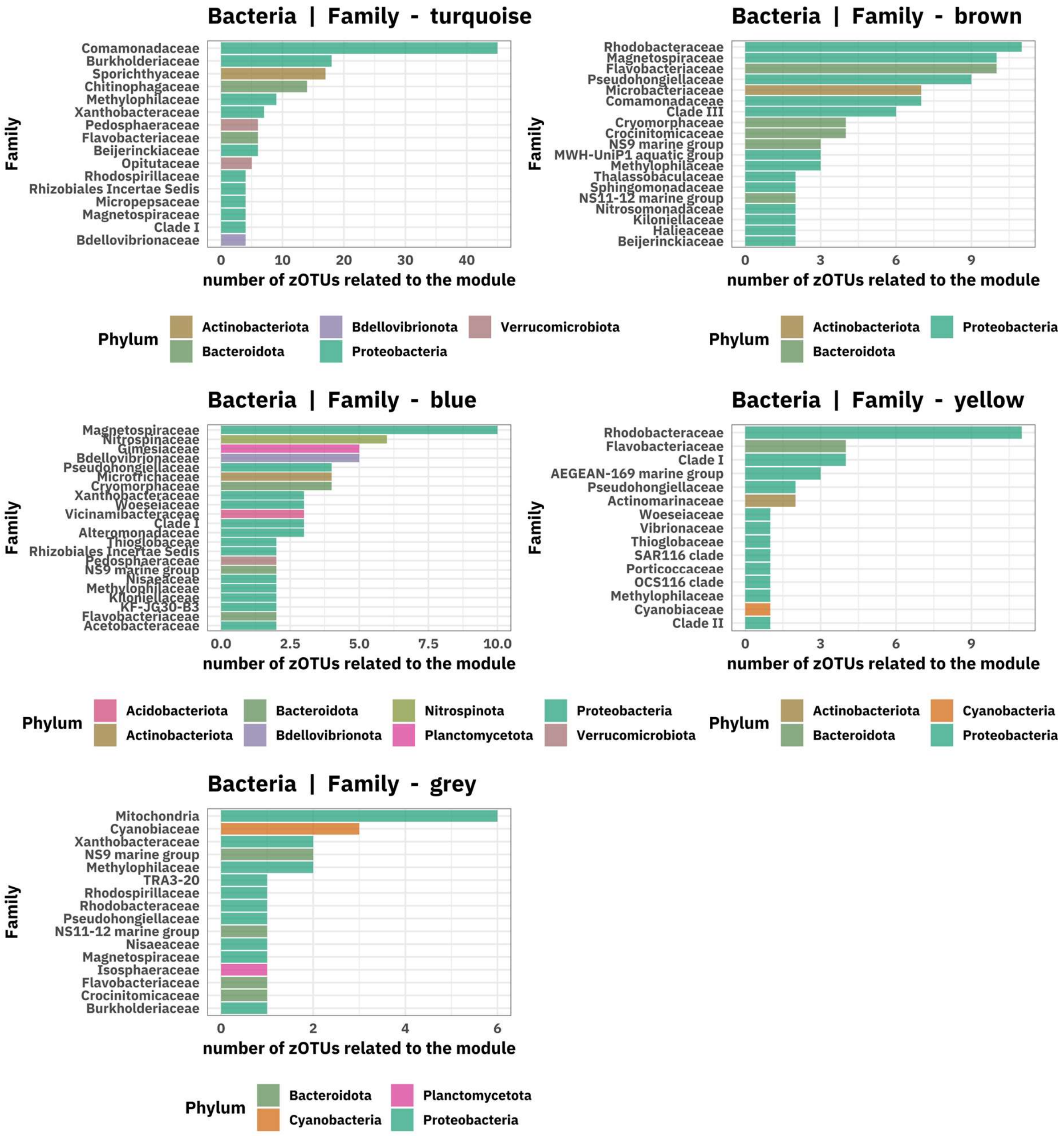


Fig. S11 - Bacterial zOTU module membership at family level. Each bar indicates the most frequent zOTUs at family level in each submodule. Different color represents different phyla. Only the 15 most abundant taxa are displayed. *Mitochondria* = zOTUs related to *Rickettsiales* (*Alphaproteobacteria*)

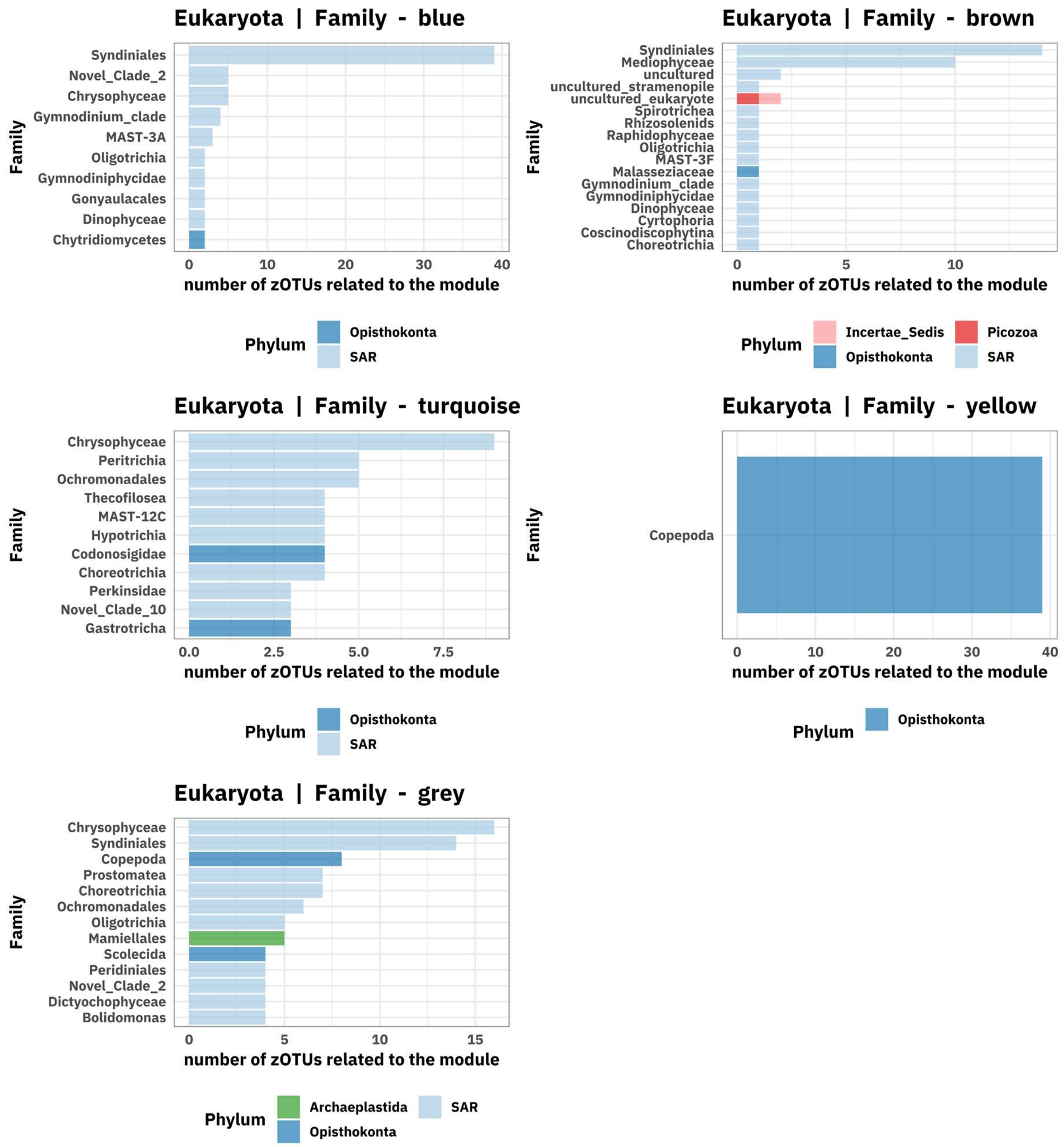


Fig. S12 - Eukaryotic zOTU module membership at family level. Each bar indicates the most frequent zOTUs at family level in each submodule. Different color represents different phyla. Only the 10 most abundant taxa are displayed.

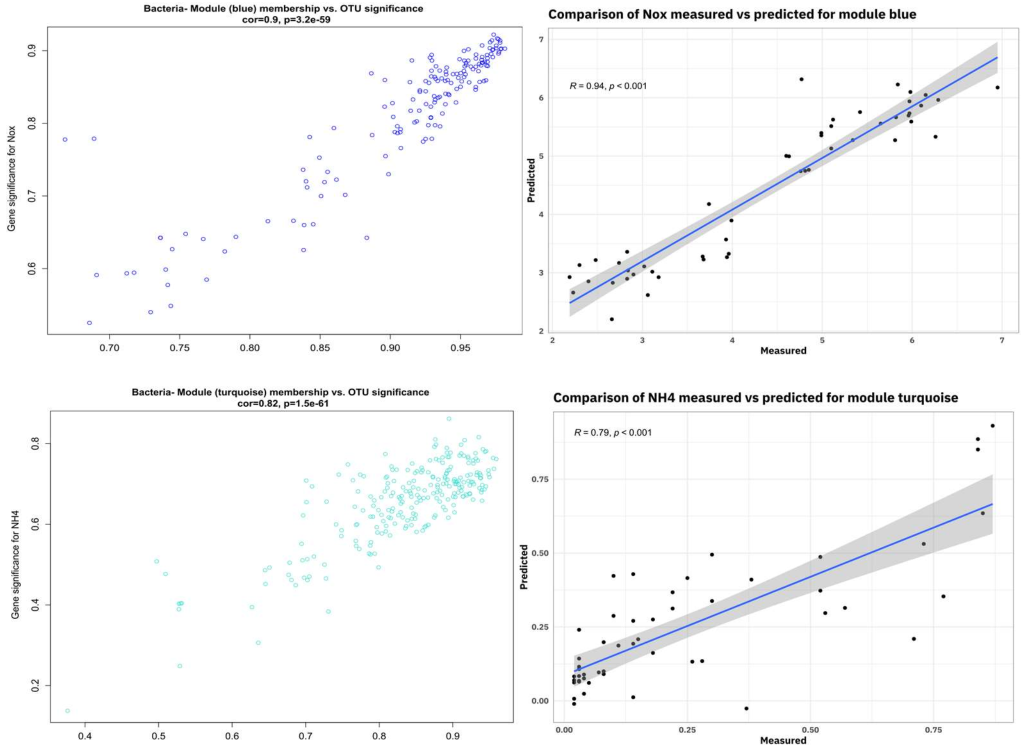


Fig. S13 - In the first column scatterplots showing bacterial zOTU significance (individual correlation to the parameter of interest) in the y-axis versus submodule membership (i.e., submodule structure: strength of an zOTU measured by the number of connections, to other zOTUs within the module) in the x-axis for the blue and turquoise submodules. Therefore, the graphs show how each taxon (each dot is an zOTU that belong to the determined module) correlated to the environmental factor of interest and how important it is to the module. Then, the zOTUs with high module membership tend to occur always connected whenever the module is represented in the environment. In the second column, Partial least squares analysis results predicting environmental variables values using CLR-transformed values of bacterial zOTUs within the module most connected to the respective environmental variable.

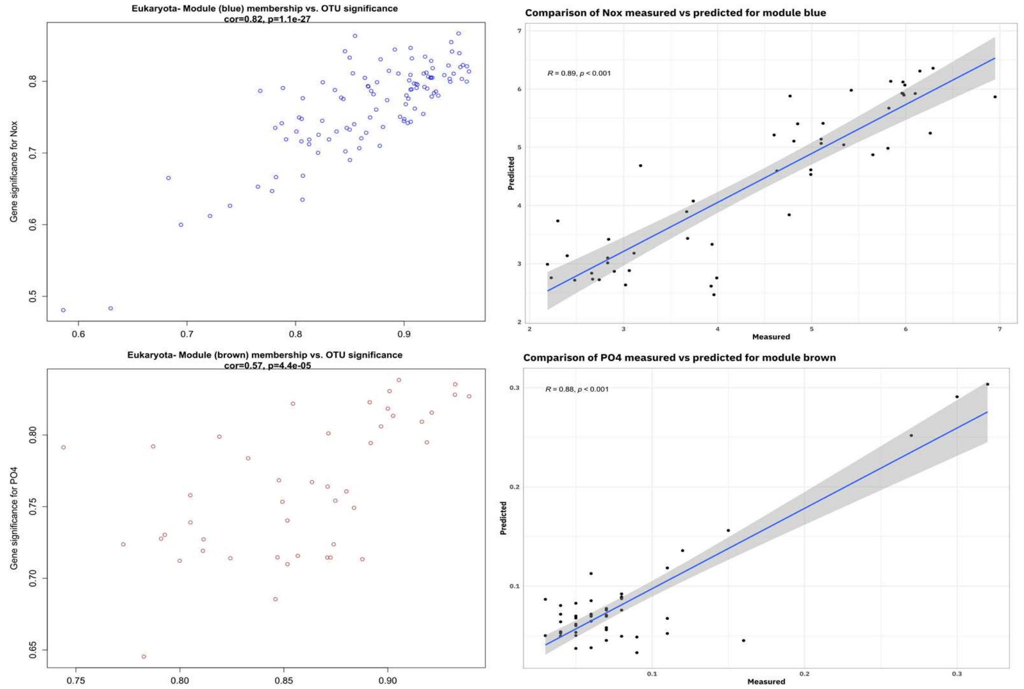


Fig. S14 - In the first column scatterplots showing eukaryotic zOTU significance (individual correlation to tto the parameter of interest) vin the y-axis versus submodule membership (submodule structure: strength of an zOTU measured by the number of connections, to other zOTUs within the module) in the x-axis for the blue and brown submodules. The graphs show how each taxon (each dot is an zOTU that belong to the determined module) correlated to the environmental factor of interest and how important it is to the module. Then, the zOTUs with high module membership tend to occur always connected whenever the module is represented in the environment. In the second column, Partial least squares analysis results predicting environmental variables values using CLR-transformed values of eukaryotic zOTUs within the module most connected to the respective environmental variable.
