## supplementary tables for "The importance of environmental parameters and mixing zone in shaping estuarine microbial communities along a freshwater-marine gradient"

**SUPPLEMENTARY MATERIAL – TABLES**

Table S1 - Environmental parameters. Sample ID can be read as the following: name of the sample and location (only for samples from the Macquarie Harbour), site (MH: Macquarie Harbour, ocean or GR: Gordon River), and water layer (S: surface or P: intermediate waters). Temp: temperature (°C), Sal: salinity (PSU), Chl: chlorophyll (μg/L), and dissolved nutrients (μM). S = surface; P = intermediate waters.

| Layer | Location | Sample ID | Temperature | Sal | Chlorophyll | $No_x$ | $PO_4$ | $SiO_4$ | $NO_2$ | $NH_4$ |
|---|---|---|---|---|---|---|---|---|---|---|
| S | A | A1_A_MH_S | 18.2 | 7.43 | 27.03 | 2.84 | 0.07 | 16.6 | 0.37 | 0.3 |
| S | A | A3_A_MH_S | 17.75 | 7.32 | 25.72 | 3.02 | 0.05 | 16.5 | 0.39 | 0.52 |
| S | A | A4_A_MH_S | 17.38 | 6.44 | 27.18 | 3.11 | 0.07 | 16.3 | 0.4 | 0.73 |
| S | B | C1_B_MH_S | 18.97 | 5.13 | 25.02 | 2.3 | 0.07 | 15.2 | 0.31 | 0.1 |
| S | B | C2_B_MH_S | 19.78 | 5.04 | 23.77 | 2.19 | 0.08 | 14.9 | 0.34 | 0.08 |
| S | B | C3_B_MH_S | 19.1 | 6.64 | 21.9 | 2.67 | 0.05 | 16.2 | 0.32 | 0.3 |
| S | B | C4_B_MH_S | 18.8 | 7.25 | 23.34 | 2.74 | 0.05 | 16.2 | 0.35 | 0.57 |
| S | B | B4_B_MH_S | 17.37 | 7.362 | 25.508 | 3.18 | 0.05 | 16.6 | 0.361 | 0.52 |
| S | B | B3_B_MH_S | 17.94 | 7.13 | 25.49 | 2.9 | 0.07 | 16.2 | 0.37 | 0.77 |
| S | B | B2_B_MH_S | 18.07 | 7.07 | 26.75 | 2.83 | 0.08 | 16 | 0.37 | 0.22 |
| S | B | B1_B_MH_S | 18.38 | 6.82 | 26.88 | 2.66 | 0.07 | 15.9 | 0.37 | 0.18 |
| S | C | D1_C_MH_S | 16.67 | 12.13 | 21.97 | 3.06 | 0.06 | 16.4 | 0.36 | 0.53 |
| S | C | D2_C_MH_S | 17.72 | 7.64 | 25.71 | 2.83 | 0.06 | 17 | 0.35 | 0.38 |
| S | C | D3_C_MH_S | 18.78 | 6.81 | 24.84 | 2.48 | 0.08 | 16.4 | 0.35 | 0.22 |
| S | C | D4_C_MH_S | 18.73 | 5.35 | 24.82 | 2.4 | 0.08 | 15.6 | 0.35 | 0.14 |
| S | C | D5_C_MH_S | 19.75 | 5.05 | 21.81 | 2.23 | 0.07 | 15.7 | 0.34 | 0.08 |
| S | GR | GR_1_S | 12.58 | 0.46 | 26.46 | 3.93 | 0.06 | 3.5 | 0.44 | 0.85 |
| S | GR | GR_2_S | 12.3 | 0.22 | 26.92 | 3.94 | 0.06 | 2 | 0.42 | 0.84 |
| S | GR | GR_3_S | 12.67 | 0.17 | 26.8 | 3.96 | 0.06 | 1.9 | 0.37 | 0.84 |
| S | GR | GR_4_S | 11.71 | 0.03 | 26.91 | 3.99 | 0.05 | 1.2 | 0.31 | 0.87 |
| S | ocean | ocean_1_S | 15.09 | 28.17 | 12.45 | 3.68 | 0.27 | 6.7 | 0.45 | 0.18 |
| S | ocean | ocean_2_S | 14.89 | 30.88 | 11.58 | 3.74 | 0.32 | 4.4 | 0.46 | 0.15 |
| S | ocean | ocean_3_S | 15.11 | 30.55 | 13.69 | 3.67 | 0.3 | 5 | 0.44 | 0.14 |
| P | A | T10_A_MH_P | 15.34 | 18.86 | 1.01 | 5.65 | 0.04 | 14.2 | 0.29 | 0.03 |
| P | A | T11_A_MH_P | 15.45 | 14.84 | 1.3 | 4.81 | 0.04 | 15.2 | 0.37 | 0.14 |
| P | A | T12_A_MH_P | 15.35 | 13.09 | 0 | 4.6 | 0.03 | 15.3 | 0.38 | 0.25 |
| P | A | T13_A_MH_P | 15.05 | 13.39 | 1.3 | 4.85 | 0.04 | 16.6 | 0.38 | 0.1 |
| P | A | A1_P_A_MH_P | 16.12 | 11.05 | 1.19 | 4.99 | 0.04 | 17.9 | 0.34 | 0.26 |
| P | A | A3_P_A_MH_P | 15.07 | 22.34 | 0 | 5.98 | 0.04 | 14 | 0.18 | 0.04 |
| P | A | A4_P_A_MH_P | 14.97 | 22.91 | 0 | 6.15 | 0.03 | 14 | 0.13 | 0.03 |
| P | B | T5_B_MH_P | 15.31 | 22.71 | 0.86 | 6.26 | 0.07 | 12.4 | 0.34 | 0.07 |

| | | | | | | | | | | |
|---|---|---|---|---|---|---|---|---|---|---|
| P | B | T6_B_MH_P | 15.08 | 23.98 | 0.79 | 5.96 | 0.07 | 12.7 | 0.13 | 0.02 |
| P | B | T7_B_MH_P | 15.35 | 22.17 | 0.88 | 5.34 | 0.05 | 12.1 | 0.16 | 0.03 |
| P | B | T8_B_MH_P | 15.52 | 19.6 | 0.91 | 5.81 | 0.05 | 13.5 | 0.27 | 0.03 |
| P | B | T9_B_MH_P | 15.79 | 15.93 | 1.19 | 4.99 | 0.04 | 14.9 | 0.41 | 0.71 |
| P | B | C1_P_B_MH_P | 15.64 | 22.74 | 0.88 | 5.1 | 0.09 | 12.4 | 0.2 | 0.04 |
| P | B | C2_P_B_MH_P | 15.23 | 23.19 | 0.87 | 5.97 | 0.08 | 12.9 | 0.27 | 0.37 |
| P | B | C3_P_B_MH_P | 15.2 | 22.7 | 0.89 | 5.99 | 0.06 | 13.2 | 0.16 | 0.02 |
| P | B | C4_P_B_MH_P | 15.15 | 22.57 | 0.89 | 6.1 | 0.06 | 13.4 | 0.18 | 0.02 |
| P | B | B4_P_B_MH_P | 14.94 | 28.07 | 0 | 6.95 | 0.16 | 12 | 0.08 | 0.04 |
| P | B | B3_P_B_MH_P | 15.09 | 23.08 | 0 | 5.82 | 0.05 | 12.4 | 0.17 | 0.05 |
| P | B | B2_P_B_MH_P | 15.21 | 21.8 | 0 | 5.84 | 0.04 | 13.7 | 0.21 | 0.02 |
| P | B | B1_P_B_MH_P | 15.18 | 21.69 | 0 | 5.97 | 0.04 | 13.5 | 0.18 | 0.03 |
| P | C | T1_C_MH_P | 14.95 | 14.36 | 0.8 | 4.77 | 0.08 | 11 | 0.14 | 0.03 |
| P | C | T2_C_MH_P | 15.26 | 22.28 | 1 | 5.1 | 0.11 | 12.5 | 0.36 | 0.11 |
| P | C | T3_C_MH_P | 15.04 | 24.72 | 0.86 | 5.42 | 0.11 | 11.2 | 0.29 | 0.03 |
| P | C | T4_C_MH_P | 15.02 | 24.9 | 0.86 | 6.29 | 0.09 | 11.6 | 0.1 | 0.02 |
| P | C | D3_P_C_MH_P | 15.6 | 22.66 | 0.88 | 4.63 | 0.15 | 12 | 0.39 | 0.28 |
| P | C | D4_P_C_MH_P | 15.81 | 23.33 | 0.85 | 4.76 | 0.12 | 12.1 | 0.29 | 0.14 |
| P | C | D5_P_C_MH_P | 15.66 | 23.06 | 0.88 | 5.12 | 0.11 | 11.9 | 0.31 | 0.08 |

Table S2 – perMANOVA and pairwise post hoc results comparing differences in microbial composition between locations and water layers. Post hoc analyses were unable to detect significant variation between the river and ocean samples.

| **Kingdom** | **Archaeal** | | **Bacterial** | | **Eukaryotes** | |
|---|---|---|---|---|---|---|
| **contrasts** | **adj.p.value** | **adonis.R2** | **adj.p.value** | **adonis.R2** | **adj.p.value** | **adonis.R2** |
| **MH_P vs GR_S** | < 0.01 | 0.35 | < 0.01 | 0.75 | < 0.01 | 0.43 |
| **MH_S vs GR_S** | 0.53 | 0.09 | < 0.01 | 0.64 | < 0.01 | 0.47 |
| **ocean_S vs GR_S** | 0.6 | 0.56 | 0.17 | 0.82 | 0.17 | 0.64 |
| **MH_S vs MH_P** | < 0.01 | 0.28 | < 0.01 | 0.61 | < 0.01 | 0.49 |
| **ocean_S vs MH_P** | < 0.01 | 0.4 | < 0.01 | 0.42 | < 0.01 | 0.29 |
| **ocean_S vs MH_S** | 0.01 | 0.26 | 0.01 | 0.35 | 0.01 | 0.43 |
| **PERMANOVA** | < 0.01 | 0.41 | < 0.01 | 0.75 | < 0.01 | 0.59 |
| **PERM_BETAD** | < 0.01 | | 0.01 | | 0.02 | |

Table S3 – Environmental correlations with db-RDA structure based on microbial compositions of all sites and water layers. NA means lack of significance ($p < 0.01$)

| **Kingdom** | **Archaea** | **Bacteria** | **Eukaryotes** |
|---|---|---|---|

| env factor | p.value ordistep | p.value env.fit | r | p.value ordistep | p.value env.fit | r | p.value ordistep | p.value env.fit | r |
|---|---|---|---|---|---|---|---|---|---|
| Chlorophyll | 0.005 | NA | 0.09 | 0.005 | 0.004 | 0.17 | 0.005 | 0.001 | 0.82 |
| PO4 | 0.005 | NA | 0 | NA | NA | 0.02 | 0.005 | 0.001 | 0.47 |
| Temperature | 0.01 | 0.007 | 0.2 | 0.005 | 0.001 | 0.57 | 0.005 | NA | 0.15 |
| SiO4 | NA | 0.001 | 0.37 | 0.005 | 0.001 | 0.45 | 0.005 | 0.001 | 0.32 |
| NH4 | NA | 0.009 | 0.19 | NA | 0.002 | 0.28 | NA | 0.001 | 0.39 |
| Sal | NA | NA | 0.15 | 0.005 | NA | 0.14 | NA | 0.001 | 0.73 |
| Nox | NA | NA | 0.05 | 0.005 | 0.004 | 0.19 | 0.005 | 0.001 | 0.62 |

Table S4 - Summary of the top 20 bacterial and eukayotic communities (agglomerated at the family level) correlations to the variable of interest found in each submodule. (n.OTUs = number of zTOUs; mean.cor = mean correlation; mean.VIP = mean VIP score)

| Submodule/ env trait | Kingdom | Class | Order | Family | nOTUs | mean.cor | mean.VIP |
|---|---|---|---|---|---|---|---|
| Blue/Nox | Bacteria | Dehalococcoidia | SAR202 clade | unknown | 4 | 0.911 | 1.200 |
| Blue/Nox | Bacteria | Nitrospinia | Nitrospinales | Nitrospinaceae | 3 | 0.910 | 1.083 |
| Blue/Nox | Bacteria | Alphaproteobacteria | Defluviicoccales | unknown | 2 | 0.900 | 1.049 |
| Blue/Nox | Bacteria | Alphaproteobacteria | Rhodospirillales | Magnetospiraceae | 2 | 0.902 | 1.085 |
| Blue/Nox | Bacteria | Alphaproteobacteria | AT-s3-44 | unknown | 1 | 0.903 | 1.033 |
| Blue/Nox | Bacteria | Alphaproteobacteria | SAR11 clade | Clade I | 1 | 0.907 | 0.911 |
| Blue/Nox | Bacteria | Babeliae | Babeliales | unknown | 1 | 0.922 | 1.065 |
| Blue/Nox | Bacteria | Bacteroidia | Flavobacteriales | Cryomorphaceae | 1 | 0.911 | 1.171 |
| Blue/Nox | Bacteria | Bacteroidia | Flavobacteriales | NS9 marine group | 1 | 0.902 | 1.261 |
| Blue/Nox | Bacteria | BD2-11 terrestrial group | unknown | unknown | 1 | 0.896 | 1.111 |
| Blue/Nox | Bacteria | Gammaproteobacteria | Thiomicrospirales | Thioglobaceae | 1 | 0.902 | 0.972 |
| Blue/Nox | Bacteria | Gracilibacteria | Candidatus Peribacteria | unknown | 1 | 0.907 | 1.101 |
| Blue/Nox | Bacteria | Subgroup 5 | unknown | unknown | 1 | 0.902 | 1.209 |
| Blue/Nox | Eukaryota | Alveolata | Protalveolata | Syndiniales | 7 | 0.836 | 1.180 |
| Blue/Nox | Eukaryota | Alveolata | Dinoflagellata | unknown | 2 | 0.822 | 1.034 |
| Blue/Nox | Eukaryota | Rhizaria | Cercozoa | Novel_Clade_2 | 2 | 0.820 | 1.042 |
| Blue/Nox | Eukaryota | Alveolata | Dinoflagellata | Gonyaulacales | 1 | 0.820 | 1.146 |
| Blue/Nox | Eukaryota | Alveolata | Protalveolata | unknown | 1 | 0.847 | 1.050 |
| Blue/Nox | Eukaryota | Chloroplastida | Charophyta | Spermatophyta | 1 | 0.855 | 1.054 |
| Blue/Nox | Eukaryota | Nucletmycea | Fungi | Chytridiomycetes | 1 | 0.842 | 1.325 |
| Blue/Nox | Eukaryota | Stramenopiles | MAST-3 | MAST-3A | 1 | 0.821 | 1.169 |
| Blue/Nox | Eukaryota | Stramenopiles | MAST-4 | unknown | 1 | 0.863 | 1.275 |
| Blue/Nox | Eukaryota | Stramenopiles | Ochrophyta | Bolidomonas | 1 | 0.833 | 1.057 |
| Blue/Nox | Eukaryota | Stramenopiles | Ochrophyta | Chrysophyceae | 1 | 0.844 | 0.924 |
| Blue/Nox | Eukaryota | uncultured eukaryote | uncultured eukaryote | uncultured eukaryote | 1 | 0.831 | 0.990 |
| Turquoise/NH4 | Bacteria | Actinobacteria | Frankiales | Sporichthyaceae | 6 | 0.785 | 1.215 |
| Turquoise/NH4 | Bacteria | Bacteroidia | Chitinophagales | Chitinophagaceae | 5 | 0.801 | 1.292 |
| Turquoise/NH4 | Bacteria | Gammaproteobacteria | Burkholderiales | Burkholderiaceae | 4 | 0.792 | 1.316 |
| Turquoise/NH4 | Bacteria | Alphaproteobacteria | Reyranellales | Reyranellaceae | 1 | 0.792 | 1.234 |

| Submodule/ env trait | Kingdom | Class | Order | Family | nOTUs | mean.cor | mean.VIP |
|---|---|---|---|---|---|---|---|
| Turquoise/NH4 | Bacteria | Bacteroidia | Chitinophagales | unknown | 1 | 0.780 | 1.162 |
| Turquoise/NH4 | Bacteria | Bacteroidia | Sphingobacteriales | Sphingobacteriaceae | 1 | 0.767 | 1.090 |
| Turquoise/NH4 | Bacteria | Gammaproteobacteria | Burkholderiales | Methylophilaceae | 1 | 0.778 | 1.330 |
| Turquoise/NH4 | Bacteria | Verrucomicrobiae | Pedosphaerales | Pedosphaeraceae | 1 | 0.776 | 1.241 |
| Brown/PO4 | Eukaryota | Alveolata | Protalveolata | Syndiniales | 7 | 0.813 | 1.011 |
| Brown/PO4 | Eukaryota | Stramenopiles | Ochrophyta | Mediophyceae | 5 | 0.810 | 1.056 |
| Brown/PO4 | Eukaryota | Alveolata | Ciliophora | Oligotrichia | 1 | 0.813 | 0.962 |
| Brown/PO4 | Eukaryota | Alveolata | Dinoflagellata | Gymnodiniphycidae | 1 | 0.792 | 0.863 |
| Brown/PO4 | Eukaryota | Stramenopiles | Labyrinthulomycetes | uncultured | 1 | 0.806 | 0.954 |
| Brown/PO4 | Eukaryota | Stramenopiles | MAST-3 | MAST-3F | 1 | 0.816 | 0.822 |
| Brown/PO4 | Eukaryota | Stramenopiles | MAST-3 | unknown | 1 | 0.823 | 0.924 |
| Brown/PO4 | Eukaryota | Stramenopiles | Ochrophyta | Coscinodiscophytina | 1 | 0.801 | 0.964 |
| Brown/PO4 | Eukaryota | Stramenopiles | Ochrophyta | Rhizosolenids | 1 | 0.827 | 1.055 |
| Brown/PO4 | Eukaryota | unknown | unknown | unknown | 1 | 0.768 | 0.952 |

Table S5 - Summary of the top 20 bacterial and eukaryotic communities (agglomerated at the family level) VIP scores found in each submodule. (n.OTUs = number of zTOUs; mean.cor = mean correlation; mean.VIP = mean VIP score)

| Submodule/ env trait | Kingdom | Class | Order | Family | n.OTUs | mean.VIP | mean.cor |
|---|---|---|---|---|---|---|---|
| Blue/Nox | Bacteria | Cyanobacteriia | Chloroplast | unknown | 1 | 1.706 | -0.778 |
| Blue/Nox | Bacteria | Gammaproteobacteria | Methylococcales | Methylomonadaceae | 1 | 1.504 | 0.860 |
| Blue/Nox | Bacteria | Alphaproteobacteria | Rickettsiales | Mitochondria | 1 | 1.491 | -0.779 |
| Blue/Nox | Bacteria | Alphaproteobacteria | Rhodospirillales | Magnetospiraceae | 2 | 1.292 | 0.828 |
| Blue/Nox | Bacteria | Bdellovibrionia | Bdellovibrionales | Bdellovibrionaceae | 2 | 1.282 | 0.881 |
| Blue/Nox | Bacteria | Dadabacteriia | Dadabacteriales | unknown | 1 | 1.263 | 0.894 |
| Blue/Nox | Bacteria | Bacteroidia | Flavobacteriales | NS9 marine group | 1 | 1.261 | 0.902 |
| Blue/Nox | Bacteria | Dehalococcoidia | SAR202 clade | unknown | 3 | 1.259 | 0.911 |
| Blue/Nox | Bacteria | Pla3 lineage | unknown | unknown | 1 | 1.227 | 0.872 |
| Blue/Nox | Bacteria | Saccharimonadia | Saccharimonadales | LWQ8 | 1 | 1.227 | -0.549 |
| Blue/Nox | Bacteria | Gammaproteobacteria | UBA10353 marine group | unknown | 1 | 1.217 | 0.886 |
| Blue/Nox | Bacteria | Subgroup 5 | unknown | unknown | 1 | 1.209 | 0.902 |
| Blue/Nox | Bacteria | Alphaproteobacteria | AT-s3-44 | unknown | 1 | 1.177 | 0.869 |
| Blue/Nox | Bacteria | Nitrospinia | Nitrospinales | Nitrospinaceae | 1 | 1.175 | 0.916 |
| Blue/Nox | Bacteria | Bacteroidia | Flavobacteriales | Cryomorphaceae | 1 | 1.171 | 0.911 |
| Blue/Nox | Bacteria | Alphaproteobacteria | SAR11 clade | Clade III | 1 | 1.166 | -0.626 |
| Blue/Nox | Eukaryota | Alveolata | Protalveolata | Syndiniales | 8 | 1.274 | 0.822 |
| Blue/Nox | Eukaryota | Alveolata | Dinoflagellata | unknown | 2 | 1.183 | 0.767 |
| Blue/Nox | Eukaryota | Alveolata | Ciliophora | Oligotrichia | 1 | 1.288 | 0.751 |
| Blue/Nox | Eukaryota | Alveolata | Dinoflagellata | Gymnodiniphycidae | 1 | 1.215 | 0.787 |
| Blue/Nox | Eukaryota | Alveolata | Dinoflagellata | Gymnodinium_clade | 1 | 1.290 | 0.742 |
| Blue/Nox | Eukaryota | Alveolata | unknown | unknown | 1 | 1.184 | 0.748 |
| Blue/Nox | Eukaryota | Nucletmycea | Fungi | Chytridiomycetes | 1 | 1.325 | 0.842 |
| Blue/Nox | Eukaryota | Picomonadida | Picomonas_judraskeda | Picomonas_judraskeda | 1 | 1.263 | 0.811 |
| Blue/Nox | Eukaryota | Rhizaria | Cercozoa | Imbricatea | 1 | 1.408 | 0.805 |
| Blue/Nox | Eukaryota | Stramenopiles | MAST-3 | MAST-3A | 1 | 1.169 | 0.821 |
| Blue/Nox | Eukaryota | Stramenopiles | MAST-4 | unknown | 1 | 1.275 | 0.863 |
| Blue/Nox | Eukaryota | unknown | unknown | unknown | 1 | 1.171 | 0.801 |
| Turquoise/NH4 | Bacteria | Actinobacteria | Frankiales | Sporichthyaceae | 4 | 1.378 | 0.764 |

| Submodule/ env trait | Kingdom | Class | Order | Family | n.OTUs | mean.VIP | mean.cor |
|---|---|---|---|---|---|---|---|
| Turquoise/NH4 | Bacteria | Bacteroidia | Cytophagales | Spirosomaceae | 3 | 1.475 | 0.715 |
| Turquoise/NH4 | Bacteria | Bacteroidia | Chitinophagales | Chitinophagaceae | 2 | 1.664 | 0.839 |
| Turquoise/NH4 | Bacteria | Gammaproteobacteria | Burkholderiales | Burkholderiaceae | 2 | 1.578 | 0.800 |
| Turquoise/NH4 | Bacteria | Gammaproteobacteria | Burkholderiales | Comamonadaceae | 2 | 1.290 | 0.744 |
| Turquoise/NH4 | Bacteria | Alphaproteobacteria | SAR11 clade | Clade III | 1 | 1.304 | 0.734 |
| Turquoise/NH4 | Bacteria | Bacteroidia | Chitinophagales | unknown | 1 | 1.318 | 0.763 |
| Turquoise/NH4 | Bacteria | Bacteroidia | Flavobacteriales | Flavobacteriaceae | 1 | 1.610 | 0.748 |
| Turquoise/NH4 | Bacteria | Bacteroidia | Sphingobacteriales | S15-21 | 1 | 1.314 | 0.622 |
| Turquoise/NH4 | Bacteria | Gammaproteobacteria | Burkholderiales | Methylophilaceae | 1 | 1.330 | 0.778 |
| Turquoise/NH4 | Bacteria | Parcubacteria | Candidatus Adlerbacteria | unknown | 1 | 1.335 | 0.759 |
| Turquoise/NH4 | Bacteria | Planctomycetes | Planctomycetales | Gimesiaceae | 1 | 1.307 | -0.384 |
| Brown/PO4 | Eukaryota | Alveolata | Protalveolata | Syndiniales | 8 | 1.128 | 0.762 |
| Brown/PO4 | Eukaryota | Stramenopiles | Ochrophyta | Mediophyceae | 4 | 1.196 | 0.803 |
| Brown/PO4 | Eukaryota | Alveolata | Dinoflagellata | Dinophyceae | 1 | 1.231 | 0.685 |
| Brown/PO4 | Eukaryota | Alveolata | Dinoflagellata | Gymnodinium_clade | 1 | 1.299 | 0.715 |
| Brown/PO4 | Eukaryota | Alveolata | Dinoflagellata | unknown | 1 | 1.075 | 0.645 |
| Brown/PO4 | Eukaryota | Nucletmycea | Fungi | Malasseziaceae | 1 | 1.133 | 0.710 |
| Brown/PO4 | Eukaryota | Picomonadida | uncultured_eukaryote | uncultured_eukaryote | 1 | 0.988 | 0.716 |
| Brown/PO4 | Eukaryota | Stramenopiles | Labyrinthulomycetes | uncultured | 1 | 0.981 | 0.739 |
| Brown/PO4 | Eukaryota | Stramenopiles | Ochrophyta | Coscinodiscophytina | 1 | 0.964 | 0.801 |
| Brown/PO4 | Eukaryota | Stramenopiles | Ochrophyta | Rhizosolenids | 1 | 1.055 | 0.827 |

Table S6 - Summary of the top 10 bacterial and eukaryotic zOTUs connectivity (node centrality) found in each submodule. (connectivity = number of edges; GS.trait = correlation value between zOTUS and environmental parameter)

| **Submodule/ env trait** | **Kingdom** | **Phylum** | **Class** | **Order** | **Family** | **connectivity** | **GS.trait** |
|---|---|---|---|---|---|---|---|
| Blue/Nox | Bacteria | Patescibacteria | Gracilibacteria | Candidatus Peribacteria | unknown | 56 | 0.907 |
| Blue/Nox | Bacteria | Proteobacteria | Alphaproteobacteria | AT-s3-44 | unknown | 40 | 0.903 |
| Blue/Nox | Bacteria | Proteobacteria | Alphaproteobacteria | Defluviicoccales | unknown | 31 | 0.898 |
| Blue/Nox | Bacteria | Actinobacteriota | Acidimicrobiia | Microtrichales | Microtrichaceae | 30 | 0.885 |
| Blue/Nox | Bacteria | Chloroflexi | Dehalococcoidia | SAR202 clade | unknown | 26 | 0.917 |
| Blue/Nox | Bacteria | Proteobacteria | Alphaproteobacteria | Rhodospirillales | Magnetospiraceae | 24 | 0.903 |
| Blue/Nox | Bacteria | Dependentiae | Babeliae | Babeliales | unknown | 22 | 0.922 |
| Blue/Nox | Bacteria | Chloroflexi | Dehalococcoidia | SAR202 clade | unknown | 21 | 0.914 |
| Blue/Nox | Bacteria | Chloroflexi | Dehalococcoidia | SAR202 clade | unknown | 20 | 0.912 |
| Blue/Nox | Bacteria | Chloroflexi | Dehalococcoidia | SAR202 clade | unknown | 13 | 0.861 |
| Blue/Nox | Bacteria | Actinobacteriota | Acidimicrobiia | Microtrichales | Microtrichaceae | 12 | 0.860 |
| Blue/Nox | Bacteria | Marinimicrobia | SAR406 clade | unknown | unknown | 11 | 0.882 |
| Blue/Nox | Eukaryota | SAR | Alveolata | Protalveolata | Syndiniales | 9 | 0.799 |
| Blue/Nox | Eukaryota | Picozoa | Picomonadida | Picomonas_judraskeda | Picomonas_judraskeda | 7 | 0.811 |
| Blue/Nox | Eukaryota | SAR | Alveolata | Dinoflagellata | Gonyaulacales | 6 | 0.820 |
| Blue/Nox | Eukaryota | SAR | Alveolata | Protalveolata | Syndiniales | 5 | 0.803 |
| Blue/Nox | Eukaryota | SAR | Alveolata | Protalveolata | Syndiniales | 5 | 0.780 |
| Blue/Nox | Eukaryota | SAR | Stramenopiles | MAST-1 | MAST-1B | 5 | 0.792 |
| Blue/Nox | Eukaryota | SAR | Stramenopiles | unknown | unknown | 5 | 0.797 |
| Blue/Nox | Eukaryota | SAR | Stramenopiles | Ochrophyta | Chrysophyceae | 5 | 0.786 |
| Turquoise/NH4 | Bacteria | Bdellovibrionota | Bdellovibrionia | Bdellovibrionales | Bdellovibrionaceae | 53 | 0.737 |
| Turquoise/NH4 | Bacteria | Proteobacteria | Alphaproteobacteria | SAR11 clade | Clade III | 42 | 0.764 |
| Turquoise/NH4 | Bacteria | Bdellovibrionota | Oligoflexia | Oligoflexales | unknown | 41 | 0.724 |
| Turquoise/NH4 | Bacteria | Bacteroidota | Bacteroidia | Sphingobacteriales | NS11-12 marine group | 33 | 0.679 |
| Turquoise/NH4 | Bacteria | Proteobacteria | Gammaproteobacteria | Burkholderiales | Comamonadaceae | 19 | 0.694 |
| Turquoise/NH4 | Bacteria | Chloroflexi | Anaerolineae | Caldilineales | Caldilineaceae | 18 | 0.686 |
| Turquoise/NH4 | Bacteria | Proteobacteria | Gammaproteobacteria | Burkholderiales | Comamonadaceae | 18 | 0.717 |

| Submodule/ env trait | Kingdom | Phylum | Class | Order | Family | connectivity | GS.trait |
|---|---|---|---|---|---|---|---|
| Turquoise/NH4 | Bacteria | Cyanobacteria | Cyanobacteriia | Chloroplast | unknown | 16 | 0.661 |
| Turquoise/NH4 | Bacteria | Bacteroidota | Bacteroidia | Chitinophagales | Chitinophagaceae | 12 | 0.722 |
| Turquoise/NH4 | Bacteria | Proteobacteria | Gammaproteobacteria | Burkholderiales | Ferrovaceae | 12 | 0.697 |
| Turquoise/NH4 | Bacteria | Proteobacteria | Alphaproteobacteria | Micropepsales | Micropepsaceae | 11 | 0.701 |
| Turquoise/NH4 | Bacteria | Bdellovibrionota | Bdellovibrionia | Bdellovibrionales | Bdellovibrionaceae | 11 | 0.707 |
| Brown/PO4 | Eukaryota | SAR | Stramenopiles | Ochrophyta | Mediophyceae | 4 | 0.795 |
| Brown/PO4 | Eukaryota | SAR | Alveolata | Protalveolata | Syndiniales | 3 | 0.794 |
| Brown/PO4 | Eukaryota | SAR | Stramenopiles | Ochrophyta | Rhizosolenids | 3 | 0.827 |
| Brown/PO4 | Eukaryota | SAR | Stramenopiles | Ochrophyta | Mediophyceae | 3 | 0.809 |
| Brown/PO4 | Eukaryota | SAR | Alveolata | Ciliophora | Choreotrichia | 3 | 0.714 |
| Brown/PO4 | Eukaryota | SAR | Stramenopiles | Ochrophyta | Mediophyceae | 3 | 0.749 |
| Brown/PO4 | Eukaryota | SAR | Alveolata | Protalveolata | Syndiniales | 3 | 0.828 |
| Brown/PO4 | Eukaryota | SAR | Alveolata | Ciliophora | Cyrtophoria | 3 | 0.724 |
| Brown/PO4 | Eukaryota | Incertae_Sedis | Telonema | uncultured_eukaryote | uncultured_eukaryote | 3 | 0.730 |
| Brown/PO4 | Eukaryota | SAR | Stramenopiles | Ochrophyta | Mediophyceae | 3 | 0.728 |
